# *C. elegans* germ cells divide and differentiate along a folded epithelium

**DOI:** 10.1101/322487

**Authors:** Hannah S. Seidel, Tilmira A. Smith, Jessica K. Evans, Jarred Q. Stamper, Thomas G. Mast, Judith Kimble

**Affiliations:** Department of Biology, Eastern Michigan University, Ypsilanti, MI; Department of Biochemistry, University of Wisconsin-Madison and HHMI, Madison, WI

## Abstract

Knowing how stem cells and their progeny are positioned within their tissues is essential for understanding their regulation. One paradigm for stem cell regulation is the *C. elegans* germline, which is maintained by a pool of germline stem cells in the distal gonad, in a region known as the ‘progenitor zone’. The *C. elegans* germline is widely used as a stem cell model, but the cellular architecture of the progenitor zone has been unclear. Here we characterize this architecture by creating virtual 3D models of the progenitor zone in both sexes. We show that the progenitor zone in adult hermaphrodites is essentially a folded epithelium. The progenitor zone in males is not folded. Analysis of germ cell division shows that daughter cells are born side-by-side along the surface of the epithelium. Analysis of a key regulator driving differentiation, GLD-1, shows that germ cells in hermaphrodites differentiate along the path of the folded epithelium, with previously described “steps” in GLD-1 expression corresponding to germline folds. Our study provides a three-dimensional view of how *C. elegans* germ cells progress from stem cell to overt differentiation, with critical implications for regulators driving this transition.

## Introduction

Adult stem cells maintain and repair tissues throughout life. Key to their function is that many stem cells reside in specialized positions within tissues. This positioning ensures that stem cells contact local support cells and receive the regulatory signals that maintain stem cells in the self-renewing state (Scheres, 2007). This positioning can also orient stem cell divisions (Yamashita and Fuller, 2008) and provide a blueprint for daughter cells as they differentiate (Yang et al., 2017). Examples abound. Stem cells in the mammalian intestine reside in crypts and differentiate as they move along intestinal villi, away from signaling Paneth cells (van der Flier and Clevers, 2009). Mouse spermatogonial stem cells reside in the basal layer of the seminiferous tubule, where they are enwrapped by supporting Sertoli cells (Chen and Liu, 2015); these stem cells differentiate as they move towards the tubule’s lumen. Likewise, in *Drosophila*, germline stem cells are anchored to and receive signals from adjacent somatic cells and differentiate after losing contact with these cells (Fuller and Spradling, 2007; Inaba et al., 2015). In plants, stem cells in the root meristem receive signals from the neighboring quiescent center and differentiate as the quiescent center is pushed farther and farther away (Aichinger et al., 2012). Thus, in each of these tissues, daughter cells lose their stem-ness and differentiate as they move away from the specialized positions occupied by stem cells. The behavior of stem cells and their daughters is therefore strongly influenced by cell position and can only be understood in the context of tissue architecture.

The *C. elegans* germline provides a tractable model for stem cell regulation (Kimble and Seidel, 2013). This tissue is maintained by a pool of germline stem cells, located in the ‘progenitor zone’ at the distal end of the gonad (Figure 1A-B). Germline stem cells are maintained in the self-renewing state through their contact with mesenchymal distal tip cells (Figure 1A-B). This contact activates GLP-1/Notch signaling in germ cells and thereby induces germ cells to transcribe key regulators of the stem cell state (Kershner et al., 2014; Shin et al., 2017). GLP-1/Notch signaling is high in germ cells in the distal end of the progenitor zone, but falls sharply as germ cells move proximally (Lee et al., 2016). By contrast, germ cells in the proximal progenitor zone increasingly express early markers of differentiation, such as GLD-1, although these cells continue to divide mitotically (e.g. Cinquin et al., 2010; Fox and Schedl, 2015; Roy et al., 2016). Germ cells stop dividing and become overtly differentiated as they exit the progenitor zone. These patterns in the progenitor zone persist despite all germ “cells” connecting to a shared cytoplasmic core (the ‘rachis’) via intercellular bridges (‘ring channels’) (Figure 1A). This simple system has enabled fundamental discoveries regarding the stem cell niche and cell-to-cell signaling (Austin and Kimble, 1987; Kimble and White, 1981), stem cell quiescence (Seidel and Kimble, 2015), and the regulatory network balancing self-renewal and differentiation (reviewed in Kershner et al., 2013).

**Figure 1.**
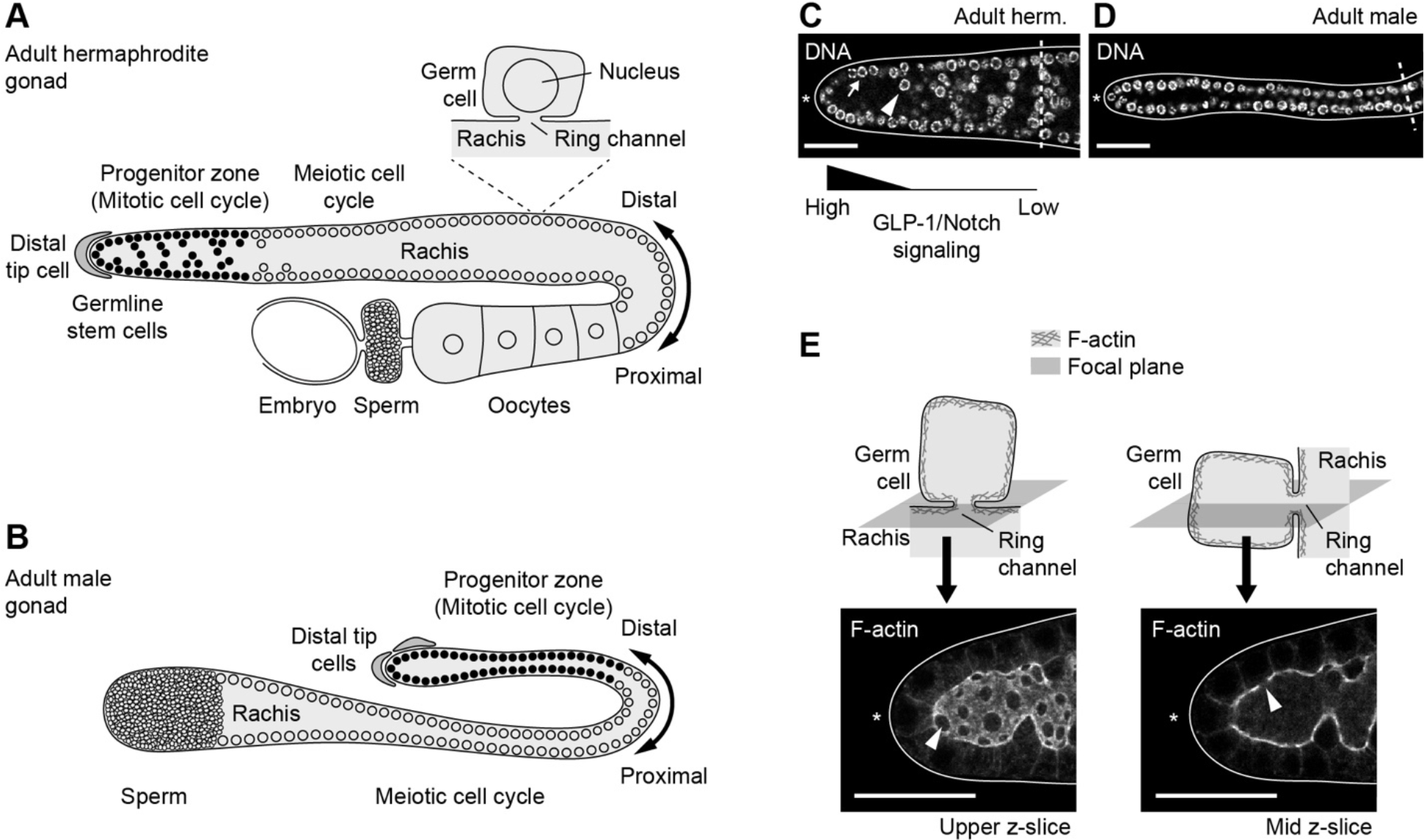
Anatomy of the *C. elegans* gonad. (A) Adult hermaphrodite gonad. (B) Adult male gonad. (C-D) Progenitor zones stained for DNA. Dashed line, boundary of progenitor zone. Arrow, example of germ cell nucleus on the outer surface of the progenitor zone. Arrowhead, example of germ cell nucleus in the interior of the progenitor zone. (E) Ring channels visualized by F-actin. Top, orientation of focal plane relative to ring channel. Bottom, adult hermaphrodite progenitor zone stained for F-actin. Arrowhead, ring channel. (C-E) Scale bar, 20 μm.

Despite intensive use of the *C. elegans* germline as a stem cell model, two key features of the progenitor zone remain poorly understood. First is the observation that some germ cell nuclei in the hermaphrodite progenitor zone reside in the interior of germline tissue, perhaps even in the middle of the rachis (Morgan et al 2010; Figure 1C, arrowhead). Other germ cell nuclei in hermaphrodites reside on the surface of the germline, their ring channels thought to point inward towards the rachis (Figure 1C, arrow). Germ cell nuclei in males virtually always reside on the surface of the germline, not in the interior (Figure 1D). How can germ cells reside in the interior of the hermaphrodite progenitor zone? How do these germ cells connect to the rachis? How does their position influence the rachis’s shape? Answering these questions is essential for understanding how the position of each germ cell relates to patterns of GLP-1/Notch signaling and to germ cell differentiation.

A second poorly understood feature of the progenitor zone concerns mitotic germ cell division. Germ cells in the progenitor zone divide continuously and asynchronously under replete conditions (Crittenden et al., 2006; Fox et al., 2011; Seidel and Kimble, 2015). Their divisions are oriented in all directions relative to the distal-to-proximal axis of the progenitor zone (Crittenden et al., 2006; Morgan et al., 2010). But how do germ cells undergo cytokinesis while remaining connected to the rachis? How is cytokinesis oriented relative to the ring channel? How can neighboring germ cells enter mitosis at different times, despite their shared cytoplasm? Answering these questions is essential for understanding cell-cycle control in the progenitor zone and lineage relationships among germ cells.

Here we characterize the cellular architecture of the *C. elegans* progenitor zone in both sexes. Our main finding is that the progenitor zone in adult hermaphrodites is a folded epithelium. The progenitor zone in males is not folded. We characterize how the hermaphrodite progenitor zone becomes folded during development, how germline folds change over time, and how these folds relate to patterns of germ cell differentiation. We also characterize mitotic germ cell divisions, showing how germ cells remain connected to the rachis during cytokinesis. Our findings provide new insights into *C. elegans* germ cell differentiation and a foundational knowledge of tissue architecture in this important stem cell model.

## Materials and methods

### Strains

N2, JK4472 *qIs154 [lag-2p::MYR::tdTomato + ttx-3p::GFP] V* (Byrd et al., 2014), NK246 *unc-119(ed4) III; qyIs8[lam-1p::lam-1::GFP + unc-119(+)] V* (Ihara et al., 2011), NK364 *unc-119(ed4) III; qyIs46[emb-9p::emb-9::mCherry + unc-119(+)] X* (Ihara et al., 2011), JK5681 *ozIs5[GLD-1::GFP] I; ltIs44 [pie-1 p::mCherry::PH(PLC1delta1) + unc-119(+)] V* (Brenner and Schedl, 2016; Kachur et al., 2008; Schumacher et al., 2005), JK3182 *gld-3(q730) nos-3(q650)/ mIn1[mIs14 dpy-10(e128)] II* (Eckmann et al., 2004), *C. briggsae* JU516, *C. remanei* MY28.

### Worm maintenance, synchronization, and staging

Worms were maintained at 20°C on nematode growth media spotted with *Escherichia coli* OP50. Nematode growth media contained 3g/L NaCl, 2.5 g/L peptone, 20g/L agar, 25 ml/L 1 M potassium phosphate buffer (1 M K_2_HPO_4_ mixed with 1 M KH_2_PO_4_ to reach a pH of 6.0), 1 mM CaCl_2_, 1 mM MgSO_4_, 5 *μ*g/ml cholesterol, and (sometimes) 2 *μ*g/ml uracil.

Animals described as ‘adult’ were staged 24 hrs post mid-L4, unless otherwise noted. Larvae were staged by the extent of gonad migration towards the vulva. Animals were classified as ‘early L4’ when gonads had migrated ~1/4 of way to the vulva; ‘mid-L4’ when gonads had migrated ~1/2-3/4 of the way to the vulva; and ‘late L4’ when gonads had migrated >3/4 of the way to the vulva. Animals were classified as ‘newly molted adults’ when animals had molted into adulthood but not yet produced embryos.

### Antibody, F-actin, and DNA staining

*C. elegans* gonads were dissected in PBSTween (PBS/0.1% Tween-20) + 0.25 mM levamisole. For staining of F-actin, GLD-1, and DAO-5, gonads were fixed in 3% paraformaldehyde for 30 min, permeabilized in 0.2% Triton X-100 for 10-30 min, then blocked in 1-3% BSA for 30 min. For alpha-tubulin and LNG staining, gonads were fixed in 3% paraformaldehyde for 15 min, permeabilized in −20°C methanol for 45 min, then blocked in 3% BSA for 30 min. Incubations with primary antibodies were performed overnight at 4°C with antibodies diluted in block as follows: rabbit anti-GLD-1 (Cinquin et al., 2010), 1/100; mouse anti-DAO-5 (Hadwiger et al., 2010), 1/10; mouse anti-alpha tubulin (Sigma, #T5168), 1/200; rabbit anti-LNG (Crittenden et al., 1994), 1/10. Incubations with secondary antibodies were performed for 1-2 hrs at room temperature, using Cy-3 donkey anti-mouse (Jackson ImmunoResearch, #715-165-151), Alexa Fluor 647 donkey anti-mouse (Invitrogen Molecular Probes, #A31571), or Alexa Fluor 647 goat anti-rabbit (Invitrogen Molecular Probes, #A21245). All secondary antibodies were diluted 1/1,000 in block. F-actin was stained by adding FITC-phalloidin (Invitrogen, #F432) into the secondary antibody incubation at a 1/50 dilution. DNA was stained by mounting gonads in Vectashield containing DAPI (Vector Labs, #H-1200).

*C. briggsae* and *C. remanei* gonads were fixed and permeabilized as for F-actin staining. Gonads were treated with 20 ug/mL RNase A at 37°C, then blocked in 3% BSA for 30 min. Gonads were stained overnight at 4°C with Alexa Fluor 488-phalloidin (Invitrogen, #A12379), diluted 1/50 in block. DNA was stained with 50 *μ*g/ml propidium iodide at room temperature for 30 min. Gonads were mounted in Vectashield as above.

### Imaging

Images of antibody stained *C. elegans* gonads were obtained on a Leica SP8 laser-scanning confocal microscope, using a z-interval of 0.3 μm. Images of *C. briggsae* and *C. remanei* gonads were obtained on a Zeiss 510 laser-scanning confocal microscope, using a z-interval of 0.5 μm.

Myristoylated tdTomato in the distal tip cell was imaged in fixed gonads. Gonads were fixed and imaged as described above for F-actin staining. Other fluorescent proteins (mCherry-PLCδ^PH^, GLD-1::GFP, Laminin::GFP, and EMB-9::mCherry) were imaged in live dissected gonads. Gonads were extruded into sperm media (50 mM HEPES, 25 mM KCl, 45 mM NaCl, 1 mM MgSO_4_, 5 mM CaCl_2_, 10 mM Dextrose, pH 7.8) containing 0.01% Tween-20, 0.25 mM levamisole, and ~50 ng/ml Hoechst 33342. Gonads were imaged immediately after dissection, on a Leica SP8 laser-scanning confocal microscope. mCherry-PLCδ^PH^ and GLD-1::GFP were imaged using a z-interval of 0.5 μm. Laminin::GFP and EMB-9::mCherry were imaged using a z-interval of 2 μm.

mCherry::PLCδ^PH^ in whole animals was imaged by mounting animals on 4% agar pads in M9 containing 10 mM NaN_3_. Animals were imaged on a Zeiss Axioimager.Z1 automated microscope, equipped with a Zeiss Axiocam.MRm camera, filter set #43 HE (excitation BP 550/25, emission BP 605/70, beam splitter FT570), and light source X-Cite 120 PC. Z-stack images were acquired using a z-interval of 0.75 μm. After imaging, animals were recovered seeded OP50 plates, then imaged again 24 hrs later. Animals landing on the agar pad on day 2 in a different orientation than on day 1 were excluded from analysis.

All images were obtained at 63X magnification, except for images of whole *gld-3 nos-3* gonads, which were obtained at 40X magnification. Within each experiment, identical imaging conditions and identical brightness adjustments were used across samples. Brightness adjustments were limited to linear adjustments, unless otherwise noted. When imaging hermaphrodites, one gonadal arm was imaged per animal; thus, gonad-to-gonad variation also reflects animal-to-animal variation.

### Scanning electron microscopy

Gonads were dissected in PBSTween + 0.25 mM levamisole, and the basal lamina was removed by digestion in 40 units/ml Type 2 collagenase (Worthington, #LS004174) in PBS for 10 min at room temperature. Gonads were fixed in 3% glutaraldehyde in PBSTween for 48 hours at 4°C; washed three times in ice-cold PBSTween + 5% sucrose; post-fixed in 2% osmium tetroxide in PBS for 60 min at room temperature; and dehydrated in an ethanol series (50%, 70%, 95%, 100%, 100%, 100%). Final drying was accomplished with hexamethyldisilazane (Electron Microscopy Sciences, #16700). Gonads were mounted on aluminum stubs using double-sided carbon tape, and stubs were sputter-coated with gold (SPI Supplies, #12150). Gonads were imaged at 5 or 8 kV accelerating potential using an Amray 1820 scanning electron microscope.

### Image annotation and modeling

Models were created from z-stack images of progenitor zones stained for F-actin, DAO-5, and DNA or progenitor zones expressing mCherry-PLCδ^PH^. Germ cell nuclei were annotated as single [*x, y, z*] points at the centers of nuclei (or, for anaphase cells, at points midway between segregating chromosomes). Nuclei in telophase cells were annotated as two separate nuclei. Positions of nuclei were annotated automatically, from DAO-5 staining, using a custom macro script in *ImageJ*. The output of this script was loaded into the *ROI Manager* of *ImageJ* and inspected and corrected manually, if necessary. Corrections were mostly limited to M-phase nuclei, which were often mis-identified by the automated script because M-phase nuclei show a different pattern of DAO-5 localization than interphase nuclei. Ring channels were annotated manually as single [*x, y, z*] points at the centers of ring channels. Regions of the rachis were annotated manually as polygons fitted to splines, using command *run("Fit Spline")*.

Models were created by plotting annotations in three dimensions in *MATLAB.* Germ cells were plotted as ‘ball-and-stick’ models, with ‘balls’ representing germ cell nuclei and ‘sticks’ connecting germ cell nuclei to their corresponding ring channels. The rachis was plotted as a volumetric mesh. This mesh was created from the rachis annotation, using the function *v2m()*, from package *iso2mesh* (Fang and Boas, 2009). Parameters values for *v2m()* were *isovalues* = *0.5*, *opt* = *2*, *maxvol* = *1*, and *method* = ‘*cgalsurf*’.

### Calculating cross-sectional area of the rachis

Cross-sectional area of the rachis was calculated in *ImageJ* from the rachis annotations of progenitor zones stained for F-actin. Rachis annotations were converted to binary images (rachis = white; outside rachis = black), and pixels inside the rachis were summed in the z-direction. The resulting z-projections were computationally straightened along a segmented line drawn manually through the midline of the progenitor zone, using command *run("Straighten…")*. Average pixel intensity in the y-direction was calculated for each x-value, using command *run("Plot Profile")*. Average pixel intensity was converted to cross-sectional area, using pixel height in y-dimension and voxel depth in z-dimension. This method accounts for the re-distribution of pixels that occurs during computational straightening.

### Scoring of cell-cycle stages

Cell-cycle stages were scored using a combination of chromosome morphology, nuclear size, and DAO-5 staining, as outlined in the table below (Crittenden et al., 2017; Seidel and Kimble, 2015). Using this method, nearly all cells that stain positive for the standard M-phase marker phospho-histone H3 were recognizable as M-phase cells (H. Seidel, personal observations).

| Stage | Nuclear size | DNA | DAO-5 localization |
| --- | --- | --- | --- |
| Interphase, special case:<br>Newly born daughter cells<br>(always occur in pairs) | Very small | Partially de-condensed;<br>localized near periphery of<br>nucleus | Nucleus<br>(nucleolar) |
| Interphase | Small to<br>large | De-condensed; localized<br>near periphery of nucleus | Nucleus<br>(nucleolar) |
| Prophase | Large | Condensed or partially<br>condensed; chromosomes<br>beginning to cluster within<br>nucleus | Nucleus (less well<br>confined to<br>nucleolus) |
| Metaphase | Large | Condensed; chromosomes<br>aligned on metaphase plate | Nucleus and<br>some cytoplasm |
| Anaphase | - | Condensed; chromosomes<br>segregating | Cytoplasm |
| Telophase | Very small | Partially de-condensed;<br>chromosomes localized in<br>clusters | Nucleus and<br>cytoplasm |

### Scoring ring channels as ‘open’ versus ‘closed’

Ring channels were scored as ‘open’ if we observed a patch of cytoplasm, devoid of F-actin, in the ring-channel passageway that was bigger than typical bare patches of F-actin in the cortical F-actin mesh. Ring channels were scored as ‘closed’ if no such patch of cytoplasm was observed. The narrowest ring channels we were able to detect as ‘open’ were ~0.3 μm in diameter.

### Scoring of germline folds and progenitor zone boundaries

Progenitor zones were scored as containing germline folds if at least one germ cell in the progenitor zone was positioned within the interior of the progenitor zone, not in contact with the outer surface of the gonad. The position of the distal-most fold was scored as the mid-point of the nucleus of the distal-most germ cell not in contact with the outer surface of the gonad. Progenitor zone boundaries were drawn as cross-sectional lines distal to the distal-most overtly differentiated germ cell. Germ cells were classified as overtly differentiated if they showed a ‘crescent’ chromosome morphology (Crittenden et al., 2017).

### Mating inhibition

Males were prevented from mating by isolating them from hermaphrodites in groups of 10-20 at the adult molt. Males were maintained without hermaphrodites for ~48 hrs before dissection.

### Quantifying GLD-1 in trios of germ cells

GLD-1 levels in trios of germ cells were quantified in z-stack images of progenitor zones stained for GLD-1 and F-actin or in progenitor zones expressing GLD-1::GFP and mCherry::PLCδ^PH^. Trios of germ cells were selected according to the following criteria: (i) all three germ cells were positioned adjacent to one another in physical space; (ii) ring channels of two of the germ cells were positioned adjacent to one another along the path of the rachis; (iii) the ring channel of the third germ cell was positioned at least four germ cell diameters away from the ring channels of the other two germ cells, as measured along the path of the rachis. To control for photo-bleaching and effects of sample depth, we limited our dataset to trios of germ cells positioned in the same focal plane. We also limited our dataset to germ cells positioned within the distal-most 15 rows of germ cells, because GLD-1 levels begin to plateau towards the proximal boundary of the progenitor zone (Brenner and Schedl, 2016). For GLD-1::GFP, we further limited our dataset to germ cells positioned in the upper half of the z-stack, because GLD-1::GFP experienced substantial photo-bleaching in the lower halves of z-stacks. We did not place a distal boundary on our dataset, but because germline folds were rare in distal-most germ cells, our dataset did not include any germ cells in the distal-most three rows of cells. We permitted our dataset to include multiple trios of germ cells from the same progenitor zone, provided that cells of each trio were positioned at least four or more germ cell diameters away from germ cells in any other trio, as measured along the path of the rachis. The number of trios qualifying to be included in our dataset ranged from zero to four, per progenitor zone.

For each germ cell in each trio, we quantified GLD-1 levels in the cytoplasm. A region of interest was drawn manually around the perimeter of the germ cell in *ImageJ*, as determined by F-actin staining or mCherry::PLCδ^PH^ localization. A second region of interest was drawn around the nucleus, as determined by DNA staining. Mean pixel intensity in the cytoplasm was calculated by excluding nuclear signal from whole cell signal. Measurements were repeated for every z-slice within a z-interval of 1.5 μm, centered on the nucleus. Measurements from different z-slices were averaged to obtain a final measurement per germ cell.

### Quantifying GLD-1 in germ cells located in backwards loops

Backwards loops were defined as regions of the rachis where the distal-to-proximal path of the rachis traveled backwards (i.e. in the proximal-to-distal direction). To quantify GLD-1 levels in germ cells along backwards loops, germ cells were chosen at each of two positions along a backwards loop: the ‘start’ of the loop (more distal as measured along the path of the rachis, more proximal as measured in physical space) and the ‘end’ of the loop (more proximal as measured along the path of the rachis, more distal as measured in physical space). Germ cells were chosen within the same focal plane, to control for effects of photo-bleaching and sample depth. When possible, three germ cells were chosen at each position. In cases where three germ cells could not be found meeting the above criteria, only two germ cells were chosen at a position. GLD-1 levels were quantified in the cytoplasm of each germ cell, as described above. Measurements were averaged across germ cells within each position to obtain a final measurement for the ‘start’ position and a final measurement for the ‘end’ position.

### Quantifying GLD-1 along the path of the rachis

GLD-1 levels along the path of the rachis were quantified in z-stack images of progenitor zones expressing GLD-1::GFP and mCherry-PLCδ^PH^. Single z-slices were identified in which the path of the rachis remained in the same focal plane for at least ~25 μm, as measured along the path of the rachis. Straight or segmented lines were drawn manually through the midline of the rachis in *ImageJ*, with a line width of 25 pixels (~3.5 μm). GLD-1 levels were quantified along the line, using the command *run("Plot Profile")*. GLD-1 levels could not be quantified throughout the entire progenitor zone (along the full trajectory of the rachis), because GLD-1 ::GFP experienced substantial photo-bleaching and was therefore dimmer in lower z-slices. Anti-GLD-1 staining was also not an appropriate tool for quantifying GLD-1 levels throughout the entire progenitor zone because this staining experienced non-uniform permeation of the tissue and was therefore dimmer at more interior regions of the rachis.

### Scoring of GLD-1 steps

GLD-1 steps were scored in z-stack images of progenitor zones stained for GLD-1 and F-actin or in progenitor zones expressing GLD-1::GFP and mCherry::PLCδ^PH^. GLD-1 steps were defined as distinct changes in GLD-1 levels occurring between neighboring patches of three or more germ cells. This definition is similar to methods used previously (Cinquin et al., 2015, 2010), but is subjective because it requires interpretation of the word ‘distinct’. Despite this subjectivity, two researchers scoring GLD-1 steps independently nearly always identified GLD-1 steps in the same locations (H. Seidel, S. Crittenden, personal observations). Randomly selected examples of GLD-1 steps are shown in Figure S5.

GLD-1 steps were scored as coincident with germline folds if germ cells on either side of the step connected to regions of the rachis separated by a distance of four or more germ cell diameters, as measured along the path of the rachis. This analysis included only GLD-1 steps in the upper half of each z-stack, to reduce internal correlations in the dataset.

## Statistics

Statistical tests were performed in *R* (*cran.r-project.org*) using the function *t.test()* or *pchisq()*. For *X*^2^ goodness-of-fit tests, tails of the expected distribution were pooled to have no expected values less than 1.0, as recommended by Cochran (1954).

## Plots

Plots were generated in *MATLAB* or using the *ggplot* package (*ggplot2.org*) for *R*.

## Results

### Structure of the progenitor zone: Folded in hermaphrodites, not folded in males

To investigate the cellular architecture of the distal gonad, we imaged progenitor zones in dissected gonads and transformed our images into virtual 3D models. Models were created by annotating the positions of germ cell nuclei, ring channels, and the rachis. Nuclei were annotated by staining DNA and DAO-5, a nucleolar protein (Hadwiger et al., 2010; Korčeková et al., 2012). Ring channels and the rachis were annotated by staining filamentous actin (F-actin), which localizes to the cortex, but is absent from ring-channel passageways (Figure 1E). This modeling approach allowed us to visualize the shape of the rachis and the orientation of each germ cell relative to the rachis (Figure 2G-H).

**Figure 2.**
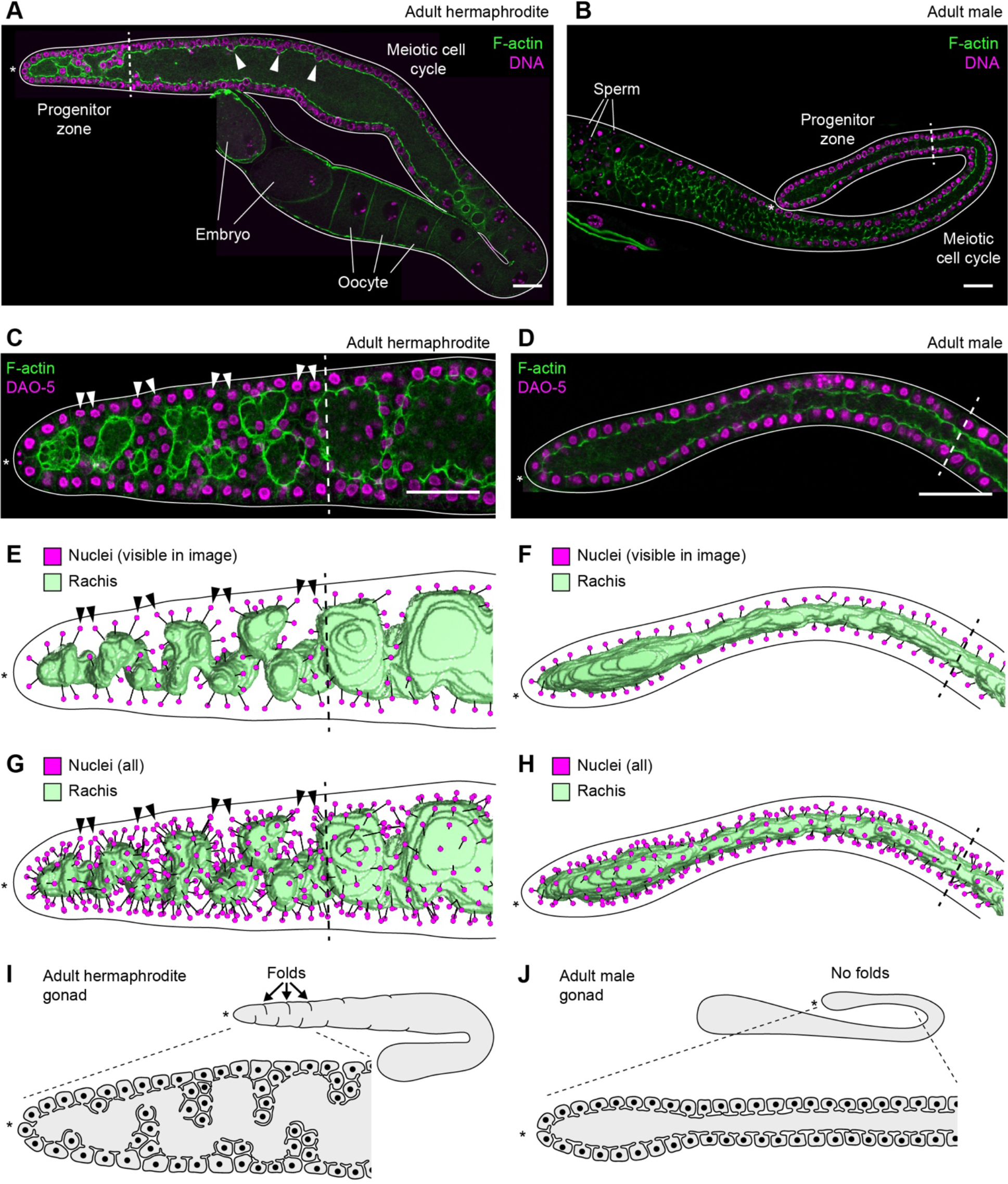
Adult progenitor zones are folded in hermaphrodites but not in males. (A-B) Adult gonads stained for F-actin and DNA. Images are composites of two or three fields of view. Arrowhead, example of shallow germline fold in the meiotic region of the gonad. (C-D) Adult progenitor zones stained for F-actin and DAO-5. Images are maximum-intensity z-projections through a z-range of 1.5 μm. Arrowhead, pair of germ cells flanking a germline fold. (A-D) Scale bar, 20 μm. (E-H) Models of rachis and germ cell nuclei. Black lines, connections between germ cell nuclei and their respective ring channels. (A-H) Solid line, outline of gonad. Dashed line, boundary of progenitor zone. Asterisk, distal end of gonad. (I-J) Schematic of progenitor-zone architecture.

Models revealed that progenitor zones in adult hermaphrodites were folded (Figure 2G). In every gonad examined (n = 41), the epithelial surface of the progenitor zone folded in and out repeatedly, bringing germ cells into the interior of the tissue. These folds caused the rachis to follow a circuitous path. As a consequence, germ cells located immediately adjacent to one another in physical space were often connected to regions of the rachis on opposite sides of a germline fold (Figure 2C and 2E, arrowheads). The placement and shape of folds varied widely from one animal to the next, except that folds were usually absent in the distal-most ~3-5 rows of germ cells (Figure S1, Figure S2). Folds became shallower and less frequent in the meiotic region of the gonad, concomitant with expansion of the rachis in this region (n = 30; Figure 2A). Folds were also visible using the plasma membrane marker mCherry::PLCδ^PH^, confirming the validity of F-actin staining (Figure S3). These results show that the progenitor zone in adult hermaphrodites is folded, and that germ cells in the interior of the progenitor zone reside within epithelial folds (Figure 2I).

We next extended our modeling to males. We predicted that male progenitor zones would lack folds, because male progenitor zones lack germ cell nuclei in the interior of the tissue (Morgan et al., 2010; Figure 1D). Consistent with this prediction, we did not observe folds in male progenitor zones (n = 35; Figure 2H). We conclude that folds are a sexually dimorphic feature of animals grown under standard laboratory conditions (Figure 2J).

### Germ cell architecture during mitotic division

To investigate mitotic germ cell divisions in cellular detail, we examined dividing germ cells in fixed tissues. We visualized the mitotic spindle with alpha tubulin and the cytokinetic ring with F-actin. F-actin also allowed us to monitor any changes to the ring channel occurring during division. This analysis revealed that germ cells divided in a stereotyped pattern in both sexes (Table 1, Figure 3). Upon entry into mitosis, germ cells closed their ring channels (Table 1, Figure 3Aii-iii, arrowhead). Ring channels transformed from an ‘open’ configuration, in which the ring-channel passageway was easily visible as a patch of cytoplasm devoid of F-actin or plasma membrane, to a ‘closed’ configuration, in which the ring-channel passageway was too narrow to be resolved by our imaging conditions (<0.3 μm in diameter; Figure 3A-B). Next, germ cells assembled the mitotic spindle parallel to the face of the rachis (n = 51 dividing cells; Figure 3C). Daughter nuclei separated along this face (Figure 3C-D). The cytokinetic ring assembled perpendicular to the face of the rachis (n = 49 dividing cells; Figure 3Avi-vii, arrow), and cleavage ingressed towards the rachis, as evidenced by late-stage (i.e. small) cytokinetic rings always abutting face of the rachis (n = 17 dividing cells; Figure 3Avii, arrow). Ring channels remained closed through the end of mitosis (Table 1), then ring channels bifurcated and re-opened in newly born daughter cells (Figure 3Aviii, arrowhead). Ring channels in daughter cells were always positioned side-by-side, flanking the site of cytokinesis (n = 50 pairs of daughter cells; Figure 3Aviii). Together, these results show that germ cells divide by closing their ring channels and cleaving along the face of the rachis (Figure 3F). An important implication of this finding is that germ cells positioned near each other along the path of the rachis are closely related by lineage; germ cells positioned near each other in physical space, by contrast, may or may not be closely related, depending on their position along the rachis (e.g. Figure 2C and 2E, arrowheads).

**Table 1.** Ring-channel configuration throughout the cell cycle.

| Cell-cycle stage | % of germ cells with ‘open’ ring-channel (n) |  |
| --- | --- | --- |
|  | Hermaphrodite | Male |
| Interphase | 98% (211) | 98% (101) |
| Prophase | 0% (76) | 0% (29) |
| Metaphase | 0% (43) | 0% (21) |
| Anaphase | 0% (25) | 0% (24) |
| Telophase | 0% (54) | 0% (29) |

**Figure 3.**
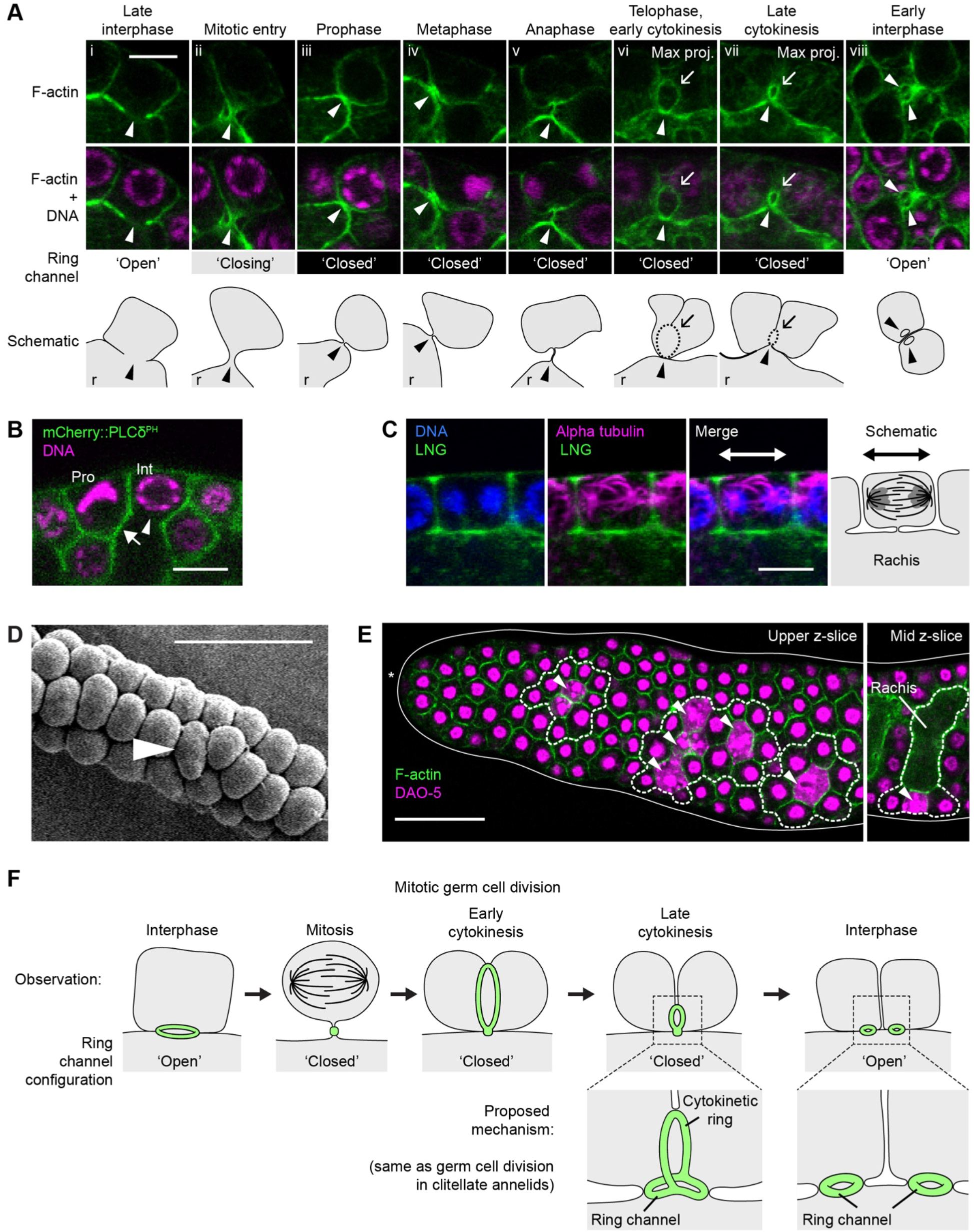
Germ cell architecture during mitotic division. (A) Germ cells stained for F-actin and DNA. Arrowhead, ring channel. Arrow, cytokinetic ring. Max proj, maximum-intensity z-projection through a z-range of 1.8 μm. r, rachis. (B) Germ cells expressing mCherry-PLCδ^PH^ and stained for DNA. Pro, prophase. Int, interphase. Arrowhead, ‘open’ ring channel. Arrow, ‘closed’ ring channel. (C) Germ cell stained for alpha-tubulin, DNA, and the LNG repeats of GLP-1 (to mark plasma membranes). (A-C) Scale bar, 5 μm. (D) Adult male progenitor zone imaged using scanning electron microscopy. Arrowhead, dividing germ cell. The face of the rachis must be immediately beneath the dividing germ cell, given that this progenitor zone comes from a male. (E) Adult hermaphrodite progenitor zone stained for F-actin and DAO-5. Left, upper z-slice (rachis not visible). Right, mid z-slice (rachis visible). Solid line, outline of gonad. Asterisk, distal end of gonad. Area enclosed by dashed line, M-phase cells, adjoining germ cells, and adjoining regions of the rachis. Arrowhead, M-phase cell. Stages of M-phase cells from left to right: telophase, anaphase, telophase, metaphase, anaphase, metaphase, anaphase. (D-E) Scale bar, 20 μm. (F) Schematic of mitotic germ cell division. The proposed mechanism is the same as germ cell division in clitellate annelids (Swiatek et al., 2009).

Our observation of ring-channel closure during division lead us to hypothesize that ring-channel closure might limit cytoplasmic exchange in or out of dividing germ cells. Consistent with this hypothesis, we observed that at least one protein—DAO-5—was seemingly unable to diffuse across ‘closed’ ring channels (Figure 3E). DAO-5 was released into germ cell cytoplasm during division, upon breakdown of the nuclear envelope, but DAO-5 did not diffuse into neighboring germ cells, nor into the adjoining region of the rachis (n = 50 dividing cells; Figure 3E). This result suggests that ‘closed’ ring channels in dividing germ cells limit the diffusion of at least some cellular contents.

### The distal tip cell extends into germline folds, but the basement membrane does not

Germ cells in the progenitor zone are regulated by their interaction with the distal tip cell, located at the distal end of the gonad (Kimble and White, 1981; Figure 1A-B). The distal tip cell in hermaphrodites forms a plexus around distal-most germ cells and extends processes proximally, some of which project deep into the interior of the progenitor zone (Byrd et al., 2014; Linden et al., 2017; Starich et al., 2014). We hypothesized that these processes reach the interior of the progenitor zone by traveling along germline folds. We imaged distal tip cells expressing myristoylated tdTomato and observed that whenever processes of the distal tip cell projected towards the interior of the progenitor zone, they always traveled along germline folds and never projected into the rachis itself (n = 17 progenitor zones; Figure 4A-B). Thus, processes of the distal tip cell in hermaphrodites extend into germline folds but not into the rachis (Figure 4D).

**Figure 4.**
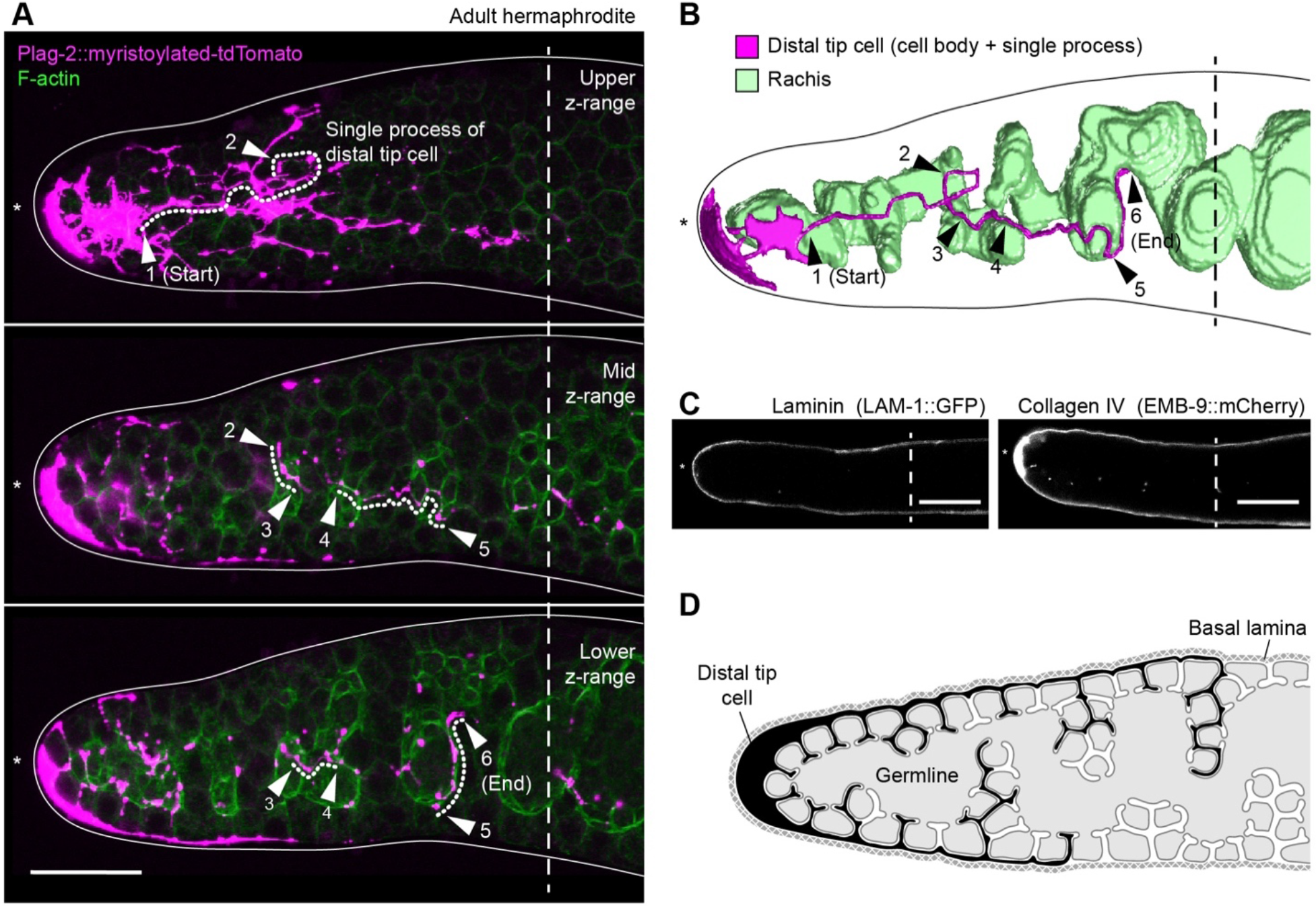
Processes of the distal tip cell extend into germline folds. (A) Adult hermaphrodite progenitor zone stained for F-actin and expressing myristoylated-tdTomato in the distal tip cell. Images are maximum-intensity z-projections through a z-range of 2.1 μm. Solid line, outline of gonad. Dotted line, single process of the distal tip cell. Arrowheads, positional markers. (B) Model of rachis and distal tip cell. (C) Adult hermaphrodite progenitor zones expressing LAM-1 ::GFP or EMB-9::mCherry. (A-C) Dashed line, boundary of progenitor zone. Asterisk, distal end of gonad. Scale bar, 20 μm. (E) Schematic of distal tip cell and basal lamina in the adult hermaphrodite progenitor zone.

A major determinant of tissue structure in animals is the basal lamina. In *C. elegans*, a basal lamina surrounds the gonad and controls gonad girth and migration of the distal tip cell (Clay and Sherwood, 2015; Kramer, 2005). We therefore asked whether the basal lamina extends into germline folds. To visualize the basal lamina, we imaged two of its components: GFP-tagged laminin and mCherry-tagged EMB-9, a type IV collagen (Ihara et al., 2011). We observed that both proteins localized to the outer surface of the hermaphrodite progenitor zone but were largely absent from progenitor zone’s interior (n = 16-20 progenitor zones; Figure 4C). We conclude that the basal lamina does not extend into germline folds (Figure 4D).

### Germline folds develop during the L4 larval stage

To understand how germline folds form during development, we examined larval germlines. The *C. elegans* germline develops from two primordial germ cells born during embryogenesis. These primordial germ cells and their descendants divide during larval development to produce an adult hermaphrodite germline containing ~1,000 germ cells per gonadal arm. We focused our analysis on the (final) L4 larval stage, because the bulk of germline expansion occurs during this stage (Kimble and Crittenden, 2005), and because germline folds appeared to be absent in larvae younger than L4. To investigate fold formation during L4, we imaged and modelled progenitor zones in four stages: Early L4s; mid L4s; late L4s; and newly molted adults. Germline folds were largely absent in early L4s but became common at later stages (Figure 5D). Moreover, folds deepened as larvae matured (Figure 5B-C). Folds in mid L4s were typically shallow, often consisting of a single germ cell tucked beneath the outer surface of the progenitor zone (Figure 5B-C). Folds in late L4s were deeper, extending farther into the interior of the progenitor zone (Figure 5B-C). Folds in newly molted adults were deeper still, similar to folds in our initial cohort of adults (aged ~24-hrs post mid-L4) (Figure 5B-C). We conclude that germline folds begin during the L4 larval stage and become more pronounced as animals reach adulthood (Figure 5A).

**Figure 5.**
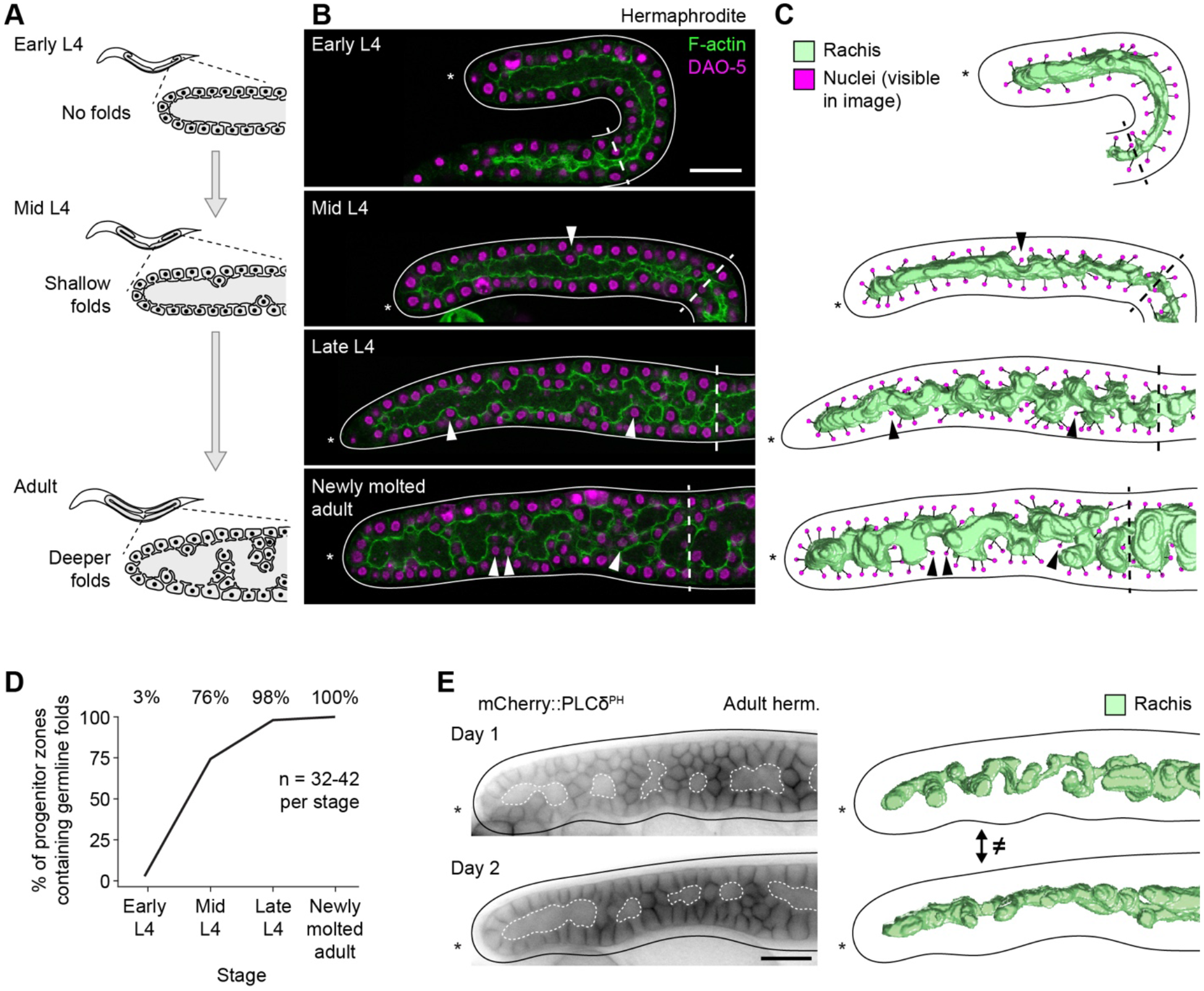
Germline folds begin during the L4 larval stage and change over time. (A) Schematic of germline development in L4 and adult hermaphrodites, summarizing findings of the current study. (B) Progenitor zones in L4 and newly molted adult hermaphrodites, stained for F-actin and DAO-5. Images are maximum-intensity z-projections through a z-range of 1.5 μm. (C) Models of rachis and germ cell nuclei. Black lines, connections between germ cell nuclei and their respective ring channels. (B-C) Arrowhead, example of germ cell positioned within a germline fold. Dashed line, boundary of progenitor zone. (D) Incidence of germline folds in L4s and newly molted adults. (E) Left, adult hermaphrodite progenitor zone expressing mCherry::PLCδ^PH^. Images have been processed with background subtraction. Dashed white line, rachis. Right, models of rachis. (B-C, E) Solid line, outline of gonad. Asterisk, distal end of gonad. Scale bar, 20 μm.

### Germline folds change over time

Germ cells in the progenitor zone are constantly dividing and moving in the distal-to-proximal direction, as cells more distal to them divide (Crittenden et al., 2006; Rosu and Cohen-Fix, 2017). We therefore hypothesized that germline folds in hermaphrodite progenitor zones might change over time, collapsing in and out, shifting location, or moving in the distal-to-proximal direction with the overall movement of cells. To test this possibility, we examined progenitor zones in live animals expressing the plasma membrane marker mCherry-PLCδ^PH^. Animals were imaged on day 1 of adulthood then again on day 2. The interval between timepoints (~24 hrs) was more than double the median cell-cycle length in the hermaphrodite progenitor zone (Fox et al., 2011; Seidel and Kimble, 2015) and should therefore allow for complete or near complete tissue turnover. In every gonad examined (n = 9), the shapes and positions of germline folds differed dramatically on the two days analyzed (Figure 5E, Figure S4). By contrast, the girth of each gonad remained similar over time (i.e. wider gonads remained wide, narrower gonads remained narrow). We conclude that germline folds are not static but instead move and change in shape over time.

### Folds are induced under conditions of germ cell crowding

What causes germline folds? Why do folds occur in hermaphrodites, but not in males? One possibility is that folds form passively, due to germ cell crowding. This possibility might explain the absence of folds in male progenitor zones, if germ cells are less crowded in males, given that sperm are produced and expelled more quickly than oocytes. To investigate this possibility, we asked whether excess germ cell crowding would induce germline folds in places where folds are normally absent. Germ cell crowding was induced in two ways. First, we prevented males from mating, which causes male gonads to become packed with sperm. Second, we used *gld-3 nos-3* loss-of-function mutants. *gld-3 nos-3* gonads fill with mitotically dividing cells (‘germline tumors’) (Eckmann et al., 2004), and hence germ cells become crowded along the length of the gonad. Both experiments resulted in germline folds: Male progenitor zones became folded in the absence of mating (n = 24; Figure 6A), and tumorous *gld-3 nos-3* gonads became folded in both sexes (n = 10-13; Figure 6B-C). The folding in *gld-3 nos-3* gonads was extensive, with the rachis transformed into a maze of narrow, convoluted passageways. Though this complexity made it difficult to map the path of the rachis in all areas of *gld-3 nos-3* gonads, we could map this path in some areas (Figure 6B‘), and folds were evident along the length of these gonads. We conclude that germline folds can form in both sexes and in all areas of the gonad, under conditions of germ cell crowding (Figure 6D-E).

**Figure 6.**
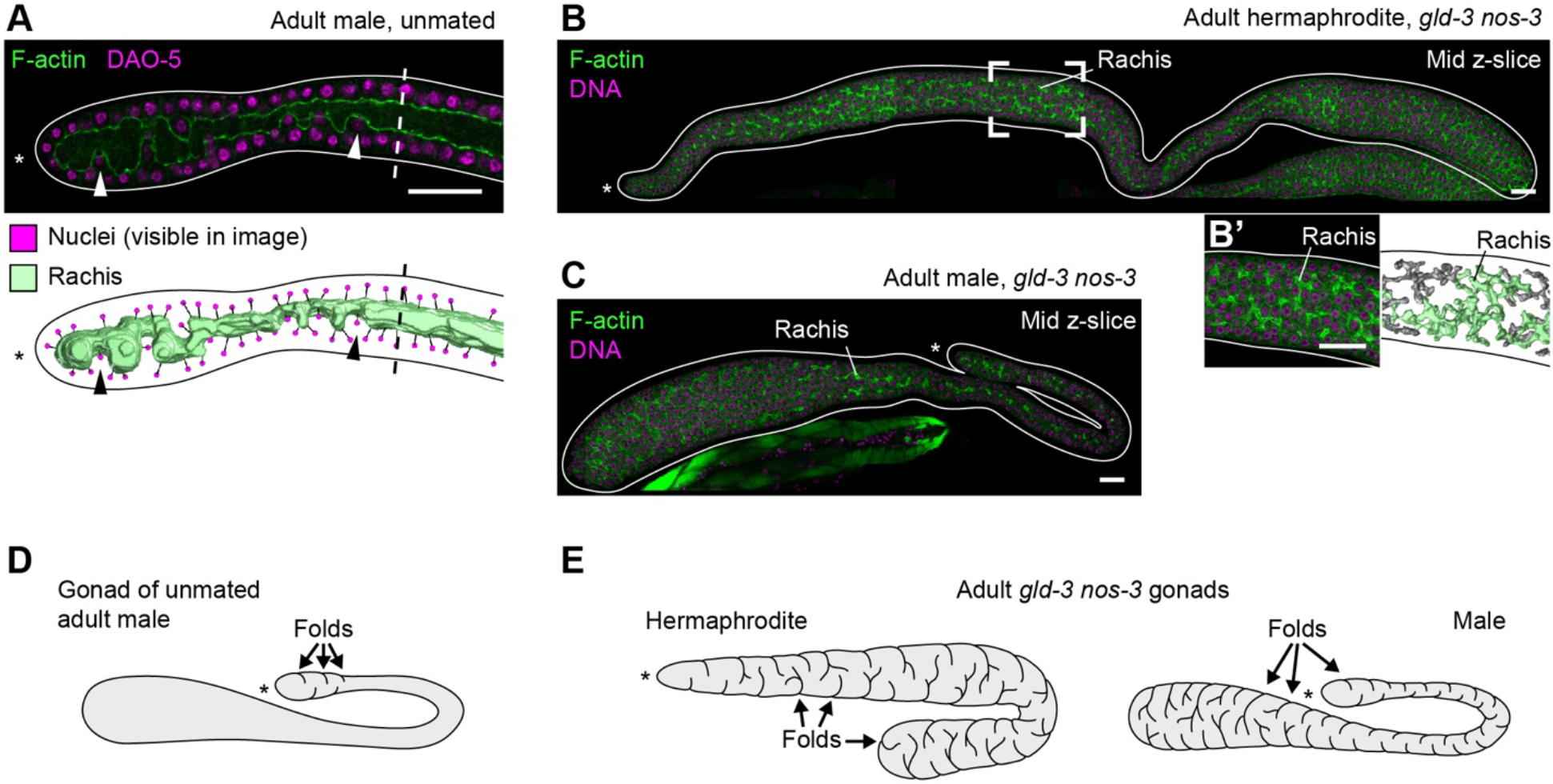
Germline folds can be induced under conditions of germ cell crowding. (A) Top, progenitor zone of unmated adult male, stained for F-actin and DAO-5. Image is maximum-intensity z-projection through a z-range of 1.5 μm. Arrowhead, example of germ cell positioned within a germline fold. Dashed line, boundary of progenitor zone. Bottom, model of rachis and germ cell nuclei. Black lines, connections between germ cell nuclei and their respective ring channels. (B-C) Gonads of *gld-3(q730) nos-3(q650)* adults stained for F-actin and DNA. Images are composites of three or four fields of view. (B’) Left, focal region marked in (B). Right, model of rachis. Green, regions of rachis contiguous within the field of view. Gray, regions of the rachis contiguous outside the field of view. (A-C) Solid line, outline of gonad. Asterisk, distal end of gonad. Scale bar, 20 μm. (D-E) Schematic of germline folds in unmated adult male and *gld-3 nos-3* adults of both sexes.

### Germ cell differentiation tracks the path of the rachis

Germ cell differentiation in *C. elegans* is traditionally viewed as occurring along a straight, distal-to-proximal path through the progenitor zone (e.g. Cinquin et al., 2010; Fox and Schedl, 2015; Lee et al., 2016). Yet our discovery of germline folds shows that germ cells do not move along a straight path, but instead move along the path defined by folds and the rachis. In addition, our analysis of germ cell division shows that lineage relationships among germ cells track the path of the rachis, not the distal-proximal axis (i.e. germ cells are mostly closely related, by lineage, to their neighbors along the rachis, not to their neighbors along the distal-proximal axis). We therefore asked if germ cell differentiation — like germ cell movement and lineage relationships — tracks the path of the rachis.

To test this possibility, we assessed the stage of differentiation in trios of germ cells positioned adjacent to one another in physical space, at essentially the same point along the distal-proximal axis. Each trio included two germ cells immediately adjacent to one another along the path of the rachis and a third germ cell positioned four or more germ cell diameters more proximally along this path (Figure 7A). The stage of differentiation of each germ cell was assessed by quantifying GLD-1, a protein whose levels rise as germ cells differentiate (Brenner and Schedl, 2016; Jones et al., 1996). GLD-1 abundance was quantified using a GLD-1 antibody (Cinquin et al., 2010) or a GLD-1::GFP transgene (Brenner and Schedl, 2016; Schumacher et al., 2005). Though this method of quantification is inherently noisy (Waters, 2009), we observed that GLD-1 levels were indistinguishable, on average, in the two germ cells positioned adjacent to one another along the rachis (Figure 7B). GLD-1 levels were two-to three-fold higher, on average, in the third germ cell (Figure 7B]). Thus, although all three germ cells were positioned at essentially the same point along the distal-proximal axis, the two germ cells lying adjacent along the rachis were at a similar stage of differentiation, whereas the third germ cell, lying several germ cell diameters more proximally along the rachis, had advanced in its differentiation. This result shows that stage of differentiation in each germ cell corresponds better with position along the rachis than with position along the distal-proximal axis of the progenitor zone.

**Figure 7.**
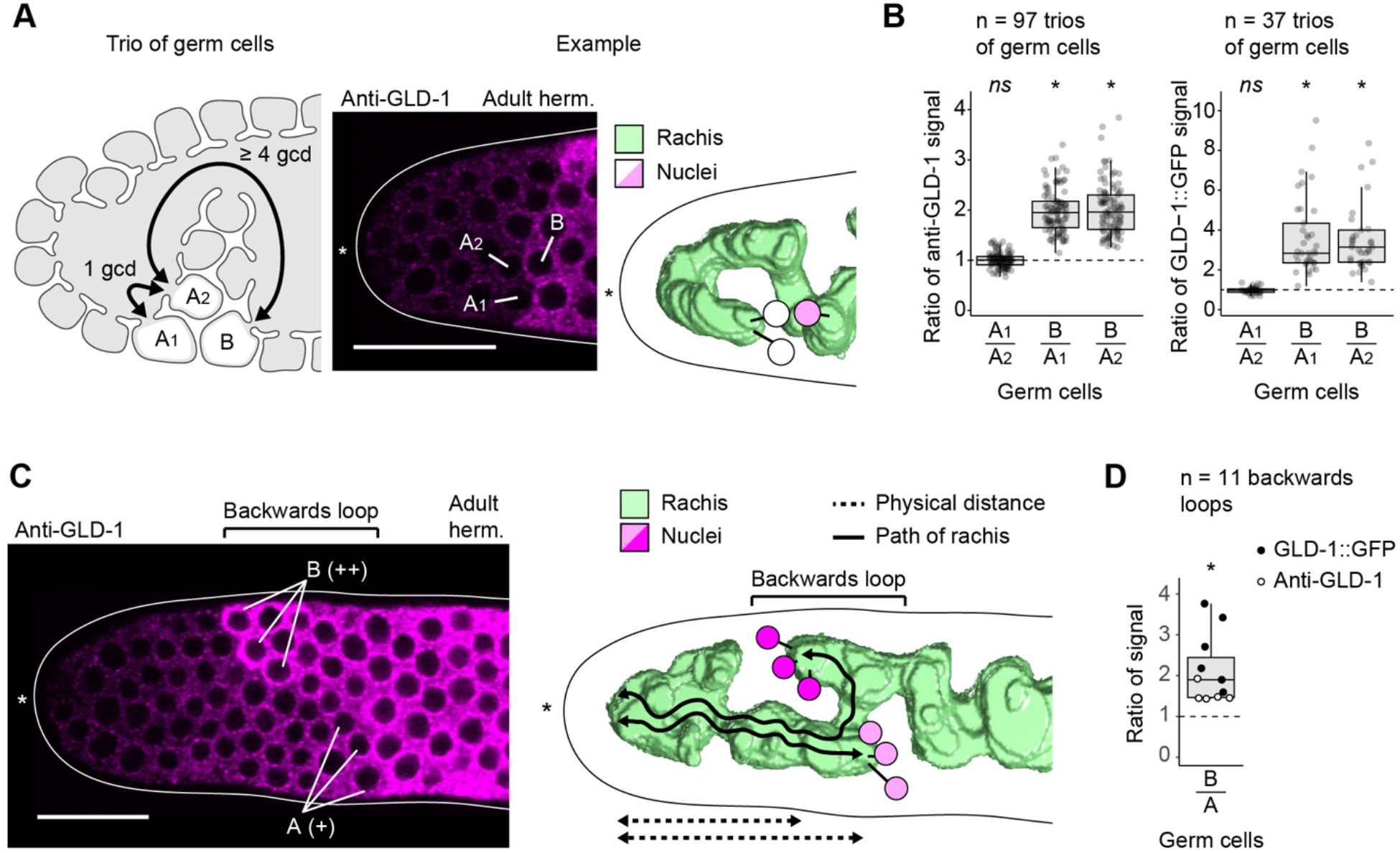
GLD-1 levels increase along the path of the rachis. (A) Left, schematic of germ cell trio used for quantification of GLD-1. gcd, germ cell diameter. Center, adult hermaphrodite progenitor zone stained for GLD-1. Right, model of rachis and germ cell nuclei. (B) Quantification of GLD-1 in germ cell trios. (C) Left, adult hermaphrodite progenitor zone stained for GLD-1 and containing a backwards loop. + and ++, relative levels of GLD-1. Right, model of rachis and germ cell nuclei. (D) Quantification of GLD-1 in germ cells along backwards loops. (A, C) Solid line, outline of gonad. Asterisk, distal end of gonad. Black lines in models, connections between germ cell nuclei and their respective ring channels. Scale bar, 20 μm. (B, D) Boxplots: center bar, median; box, interquartile range (IQR); whiskers, most extreme value within 1,5*IQR from box. dashed line, ratio of 1. *ns*, p > 0.05, paired t-test of GLD-1 levels in A_1_ versus A_2_. *, p < 0.01, paired t-test of GLD-1 levels in B versus A_1_, A_2_, or A.

As a second test of the idea that germ cell differentiation tracks the path of the rachis, we examined GLD-1 levels in the subset of progenitor zones where the rachis had looped backwards on itself. Backwards looping was observed in ~15% (n = 75) of progenitor zones and allowed us to compare GLD-1 levels among germ cells whose positions along the distal-proximal axis were reversed relative to their positions along the path of the rachis. We observed that in every backwards loop, GLD-1 levels were lower in germ cells at the start of the loop than in germ cells residing closer to the end of the loop (Figure 7C-D). This result confirms that GLD-1 levels increase as germ cells move proximally along the rachis, even when the path of the rachis deviates dramatically from a straight, distal-to-proximal trajectory through the progenitor zone. We conclude that germ cell differentiation tracks the path of the rachis, rather than a straight, distal-to-proximal path through the progenitor zone.

### Germline folds create the illusion of step-like changes in GLD-1 expression levels

GLD-1 expression in the hermaphrodite progenitor zone is often patchy — GLD-1 levels often change abruptly, in steps between neighboring groups of germ cells (Cinquin et al., 2015, 2010; Figure 8A, Figure S5). Complex models have been proposed to explain these GLD-1 steps (Cinquin et al., 2015), but our discovery of germline folds suggested a simpler idea: GLD-1 steps might be “illusions” created by germline folds bringing together, in physical space, germ cells distant along the rachis. Under this scenario, GLD-1 levels might rise gradually (not abruptly) along the path of the rachis, but this rise might appear abrupt when distant groups of germ cells are brought together by a germline fold. To test this hypothesis, we mapped GLD-1 steps relative to germline folds. GLD-1 steps were identified using a GLD-1 antibody or a GLD-1:GFP transgene, and germline folds were mapped using F-actin staining or expression of the plasma membrane marker mCherry-PLCδ^PH^. We observed that GLD-1 steps always coincided with germline folds: Germ cells on opposite sides of a GLD-1 step were always positioned on opposite sides of a germline fold (Table 2, Figure 8A). GLD-1 steps never passed through the cytoplasm of the rachis and never crossed between germ cells at the same point along the rachis. Instead, GLD-1 levels always changed gradually when measured along the rachis (n = 20; Figure 8B). These results suggest that GLD-1 levels rise gradually as germ cells differentiate, and that this rise only appears abrupt when viewed across a germline fold.

**Table 2.**
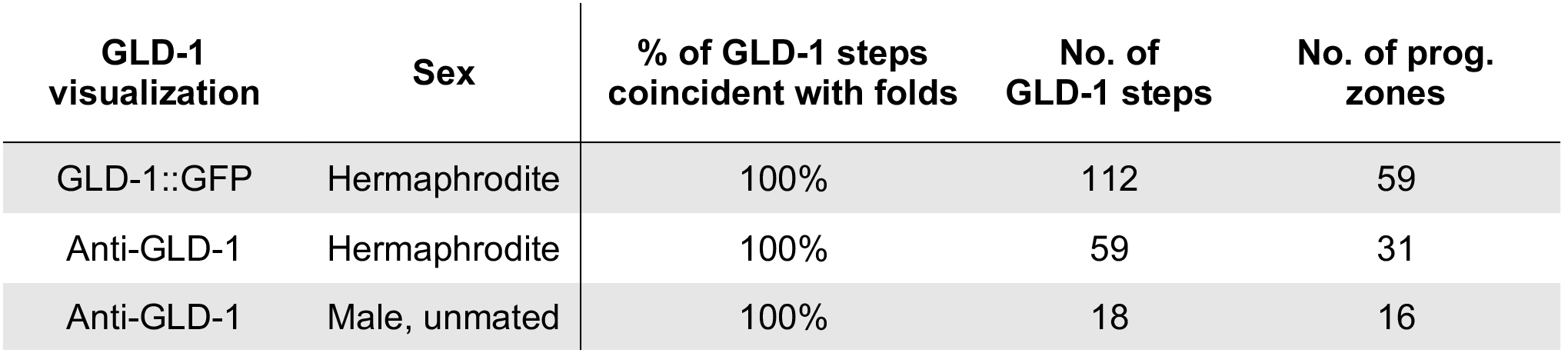
Incidence of GLD-1 steps relative to germline folds.

**Figure 8.**
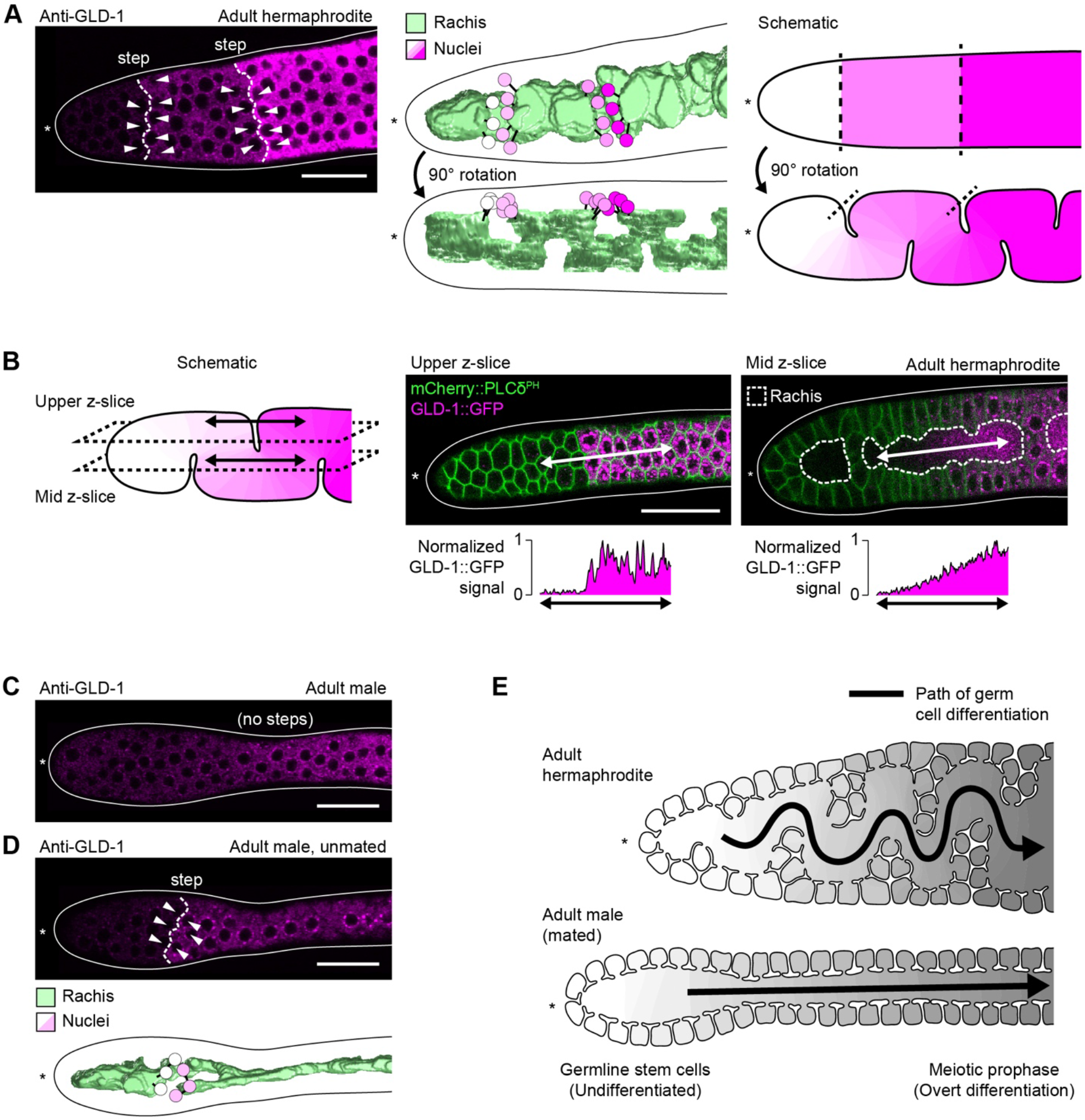
Germline folds create the illusion of abrupt changes in GLD-1 expression levels. (A) Example of GLD-1 steps. Left, adult hermaphrodite progenitor zone stained for GLD-1. Center, model of rachis and germ cell nuclei. Right, schematic of germline folds. (B) Center and right, adult hermaphrodite progenitor zone expressing GLD-1 ::GFP and mCherry::PLCδ^PH^. Bidirectional arrow, axis of GLD-1 ::GFP quantification. GLD-1 ::GFP levels were quantified across germline fold (center) and along the path of the rachis (right). Left, schematic of germline folds in the progenitor zone shown center and right. (C) Progenitor zone of adult male grown under standard laboratory conditions and stained for GLD-1. (D) Top, progenitor zone of adult male prevented from mating for two days and stained for GLD-1. Bottom, model of rachis and germ cell nuclei. (A-D) Solid line, outline of gonad. Asterisk, distal end of gonad. Scale bar, 20 μm. (A, D) Dashed line, GLD-1 step. Arrowhead, germ cell included in model. Black lines in models, connections between germ cell nuclei and their respective ring channels. (E) Model for germ cell differentiation progressing along the path of the rachis.

As a second test of our hypothesis that GLD-1 steps are an outcome of germline folds, we examined GLD-1 expression in males. Male progenitor zones normally lack GLD-1 steps (Cinquin et al., 2015; Figure 8C), and males also normally lack germline folds (Figure 2J). We predicted that if GLD-1 steps are an outcome of germline folds, then inducing germline folds in males should also induce GLD-1 steps. Consistent with this prediction, male progenitor zones with germline folds (induced by absence of mating) also developed GLD-1 steps (Table 2, Figure 8D). These GLD-1 steps in males, like GLD-1 steps in hermaphrodites, always coincided spatially with germline folds (Table 2). We conclude that GLD-1 steps can occur in both sexes, and that GLD-1 steps always coincide with germline folds.

### Germline folds are conserved in> *C. briggsae* and *C. remanei*

We asked whether germline folds were conserved in the related nematode species *C. briggsae* (hermaphrodite/male) and *C. remanei* (female/male). We observed that progenitor zones in these species were structured similarly to progenitor zones in *C. elegans*, with folds found consistently in *C. briggsae* hermaphrodites and *C. remanei* females (n = 10 per species, Figure S6). Yet in contrast to *C. elegans* males (Figure 2J), germline folds were also found in *C. briggsae* and *C. remanei* males (n = 10 per species; Figure S6). Thus, germline folds are largely conserved in the *Elegans* group of *Caenorhabditids*, although the sexual dimorphism of folding varies among species.

## Discussion

Our study describes the three-dimensional structure of the *C. elegans* progenitor zone, a key model for understanding germ cell development and stem cell control. Our main result is that the progenitor zone in adult hermaphrodites is folded. The epithelial surface of the hermaphrodite progenitor zone folds in and out, bringing germ cells into the interior of the tissue and causing the rachis to follow a circuitous, folded path (Figure 2I). The male progenitor zone, by contrast, is not folded (Figure 2J). Germ cells are born side-by-side along the path of the rachis (Figure 3F), and position along this path determines a germ cell’s stage of differentiation (Figure 8E). Germline folds in hermaphrodites begin during L4 larval development (Figure 5A), change over time (Figure 5E), and form ectopically under conditions of germ cell crowding (Figure 6E). These findings redefine spatial relationships within the progenitor zone and provide a new spatial framework for future studies in this important system.

### New view of germ cell differentiation

*C. elegans* germ cells differentiate as they move proximally, away from the distal tip cell (Kimble and Seidel, 2013). Past models measured this process as a function of germ cell position along the straight, distal-proximal axis of the progenitor zone. Past models asserted that germ cells positioned more distally within the progenitor zone would be at an earlier stage of differentiation than germ cells positioned more proximally. Likewise, germ cells positioned at the same point along the distal-proximal axis were assumed to be at similar stages of differentiation.

Our study refines this traditional model. While the general principle remains true that germ cells differentiate as they move proximally, we now find that the path of germ cell movement does not always follow a straight line. In males, the path of germ cell movement is indeed straight, consistent with the traditional view (Figure 8E). In hermaphrodites, by contrast, germ cells move proximally within germline folds. The path defined by these folds is winding and circuitous and hence does not follow a straight line (Figure 8E). Thus, germ cell differentiation in both sexes advances as germ cells move proximally along the path of the rachis, although the shape of this path in hermaphrodites is complex.

Most studies of germ cell differentiation in *C. elegans* have not considered the proximal movement of germ cells along a circuitous path. Given our results, we suggest that germ cell differentiation is driven by factors that function within germ cells as they move proximally along the rachis, rather than factors that function by tracking position along the distal-proximal axis. A simple model consistent with our study is that germ cells begin differentiating after exiting a distal-most pool of naïve germ cells, then become overtly differentiated after a fixed number of cell divisions or a fixed time interval thereafter. This model is consistent with GLP-1/Notch signaling being confined to the distal-most few rows of germ cells (Lee et al., 2016) and with germ cells differentiating after one or two cell divisions following loss of GLP-1/Notch signaling (Fox and Schedl, 2015).

Our study provides a simple explanation for apparent “steps” of gene expression in the progenitor zone that had been puzzling. The expression of many germ cell regulators is graded in the progenitor zone (e.g. Brenner and Schedl, 2016; Kershner et al., 2014; Lee et al., 2016), but expression of these regulators can be patchy, characterized by abrupt, step-like changes in expression levels between neighboring groups of germ cells. These steps are most pronounced for GLD-1 (Cinquin et al., 2010), but have also been observed for other germ cell regulators (H. Shin, K. Haupt, H. Seidel, personal observations). A previous study proposed that these GLD-1 steps reflect a slowing of distal-to-proximal diffusion at specific locations in the progenitor zone, perhaps caused by constriction of the rachis (Cinquin et al., 2015). Our clarification of progenitor-zone architecture reveals a simpler explanation. We find that GLD-1 steps correspond to germline folds. These folds bring together germ cells from different points along the rachis and hence at different stages of differentiation. We therefore suggest that the GLD-1 steps do not reflect abrupt changes in expression. We cannot exclude the possibility that germline folds influence germ cell differentiation, but we instead favor a simpler model in which germ cells differentiate along the path of the rachis, independent of the shape and placement of germline folds. This view is consistent with germline folds being highly variable in hermaphrodites (Figure S1, Figure S2), including some animals with very limited folds (Figure S1B), and with folds being absent in males (Figure 2J). This view is also consistent with our finding that germline folds do not occur at reproducible locations in the progenitor zone (Figure S2F).

Germ cell position along the distal-proximal axis is commonly used as a metric for characterizing gene expression in the progenitor zone, especially with respect to germ cell differentiation (e.g. Brenner and Schedl, 2016; Lee et al., 2016). This metric is user-friendly and will undoubtedly remain valuable, but we emphasize that position along the distal-proximal axis can be a poor proxy for a germ cell’s stage of differentiation. Germ cells residing at essentially the same point along the distal-proximal axis are sometimes at very different stages of differentiation, and the direction of differentiation along this axis is reversed, where the path of the rachis loops backwards on itself. Thus, germ cell position along the distal-proximal axis is imperfectly correlated with stage of differentiation, and future studies must take this imprecision into account.

### Germ cell division and ring-channel closure

Germ cells in virtually all animals are syncytial, connected to other germ cells via intercellular bridges (‘ring channels’ in *C. elegans*) (Greenbaum et al., 2011; Matova and Cooley, 2001). These bridges arise in many species through incomplete cytokinesis and stabilization of the cytokinetic ring (Haglund et al., 2011). Incomplete cyokinesis occurs in the *C. elegans* embryo when the P_4_ cell division gives rise to the two primordial germ cells (Goupil et al., 2017). This mechanism differs from our observations in germ cells of L4 larvae and adults, in which the ring channel bifurcates during division (Figure 3F). This latter mechanism is reminiscent of primordial germ cell formation in *Drosophila* (Cinalli and Lehmann, 2013) and is strikingly similar to germ cell division in clitellate annelids (earthworms and leeches) (Swiatek et al., 2009). Germ cells in clitellate annelids connect to a shared cytoplasmic core (Urbisz et al., 2015), and ring channels in this taxon bifurcate during division by being ‘pinched’ in two by the cytokinetic ring (as shown by transmission electron microscopy in Swiatek et al., 2009). We hypothesize that ring channels in*C. elegans* bifurcate using this same mechanism (Figure 3F), although future work is needed to image the bifurcation at higher resolution.

Ring channels in *C. elegans* germ cells close during division. One possible function of this closure is to block diffusion of cytoplasm into or out of dividing cells. Such blockage would prevent cell-cycle regulators from leaving the cell, thus allowing neighboring germ cells to cycle asynchronously, despite each cell connecting to a shared rachis cytoplasm. Such blockage is consistent with our observation that ring-channel closure blocks diffusion of DAO-5 from mitotic cells (Figure 3E). Similar blockage has been reported in *C. elegans* embryos, where the small size or composition of the intercellular bridge connecting primordial germ cells limits diffusion between them (Amini et al., 2014; Goupil et al., 2017). Ring-channel closure during division might be regulated by the same factors regulating ring-channel size in meiosis (Rehain-Bell et al., 2017) and cellularization of the oocyte (Lee et al., 2017). Ring channels are enriched for regulators of contractility (Amini et al., 2014; Maddox et al., 2005; Zhou et al., 2013) and might close via contraction of an actomyosin ring; alternatively, ring channels might close via release of a tension force holding the ring channel open during interphase.

### Fold formation and function

Many tissues form folds — brains, guts, kidneys, and lungs, to name a few. Folds can form through active cellular mechanisms (e.g. polarized contraction), passive physical mechanisms (e.g. mechanical instability) or both (Andrew and Ewald, 2010; Nelson, 2016; Pearl et al., 2017). Folds in epithelial sheets are often formed by unequal growth rates. When an epithelial sheet grows faster than its underlying substrate, the sheet will buckle and form a fold (reviewed in Nelson, 2016; Taber, 2014). We hypothesize that this same mechanism is responsible for forming folds in the *C. elegans* germline. Our model is that as germ cells divide, the cellular surface of the germline outgrows the basal lamina surrounding the gonad. As a consequence, the germline epithelium buckles inward to form a fold. This model is consistent with the shape and placement of germline folds being highly variable from one animal to the next (Figure S1, Figure S2) and with germline folds being induced under conditions of germ cell crowding (Figure 6).

Folds increase the surface area of an epithelium. This increase in the *C. elegans* germline allows the progenitor zone to package more germ cells into a space of confined dimensions. Folds likely do not serve a major role in stem cell control, because folding is absent in males and is sometimes very limited even in hermaphrodites (e.g. Figure S1B). We suggest instead that the increased number of germ cells accommodated by folds is critical for expanding the rachis to generate and supply large oocytes.

## Acknowledgements

We thank Sarah Crittenden for helpful discussions, Jane Hubbard for pointing out useful references, and David Greenstein for advice on preparing samples for scanning electron microscopy. We thank Jadwiga Forster for media preparation, Peggy Kroll-Conner for strain maintenance, and Janie Lawson, Glenn Walker, and Bob Winning for comments on the manuscript. For strains, we thank David Sherwood, Tim Schedl, and the *Caenorhabditis* Genetics Center, which is supported by NIH (P40 OD01044). JK is an investigator of the Howard Hughes Medical Institute. TAS was supported by a Symposium Undergraduate Research Fellowship Award donated by Bill Fennel.

---

**Figure S1.**
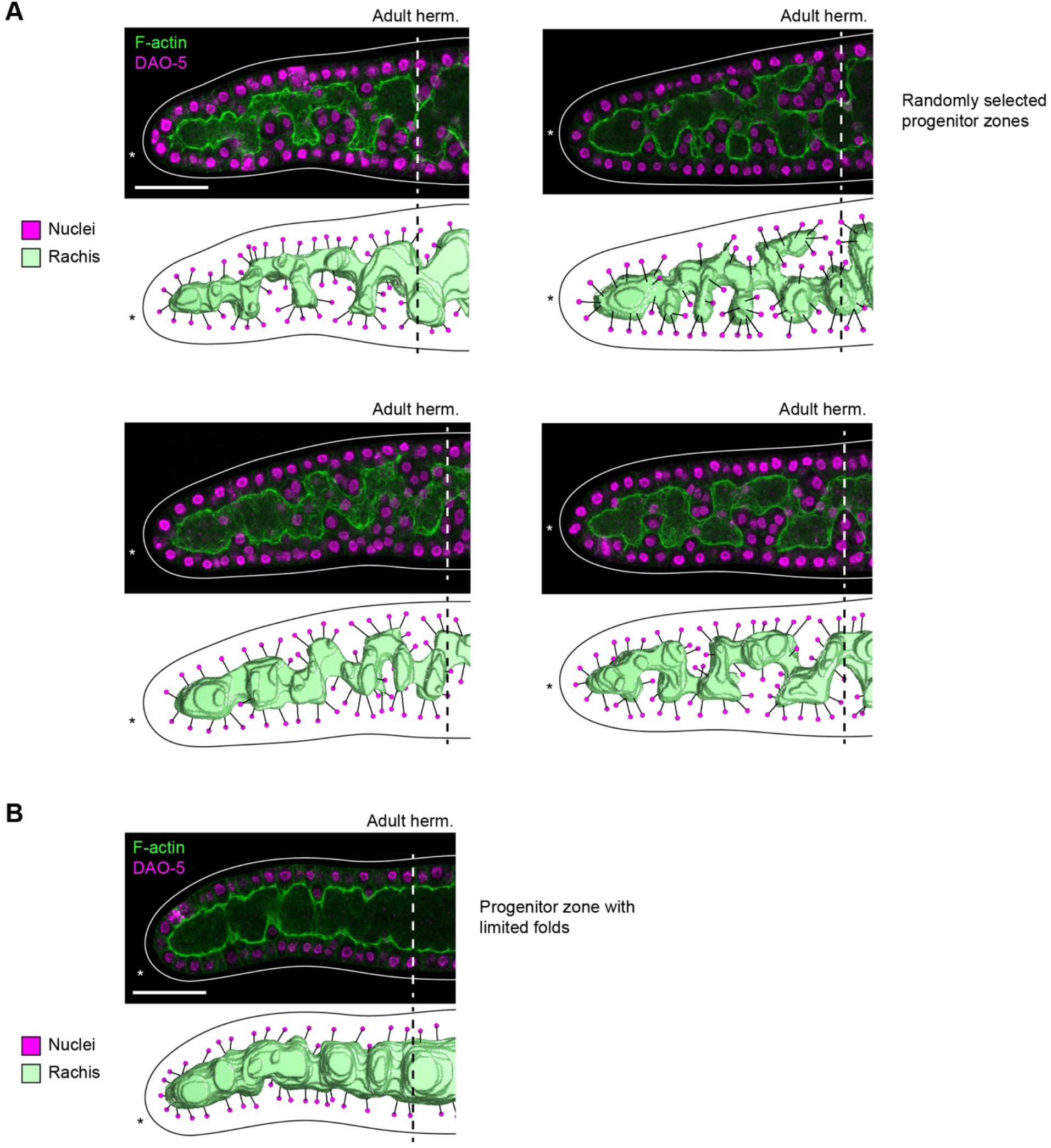
Additional examples of adult hermaphrodite progenitor zones. Top of each pair, adult hermaphrodite progenitor zone stained for F-actin and DAO-5. Bottom of each pair, model of rachis and germ cell nuclei. Images are maximum-intensity z-projections through a z-range of 1.5 μm. Models include only germ cell nuclei visible in images. Solid line, outline of gonad. Dashed line, boundary of progenitor zone. Asterisk, distal end of gonad. Black lines in models, connections between germ cell nuclei and their respective ring channels. Scale bar, 20 μm. (A) Randomly selected examples. (B) Example of progenitor zone with limited folding.

**Figure S2.**
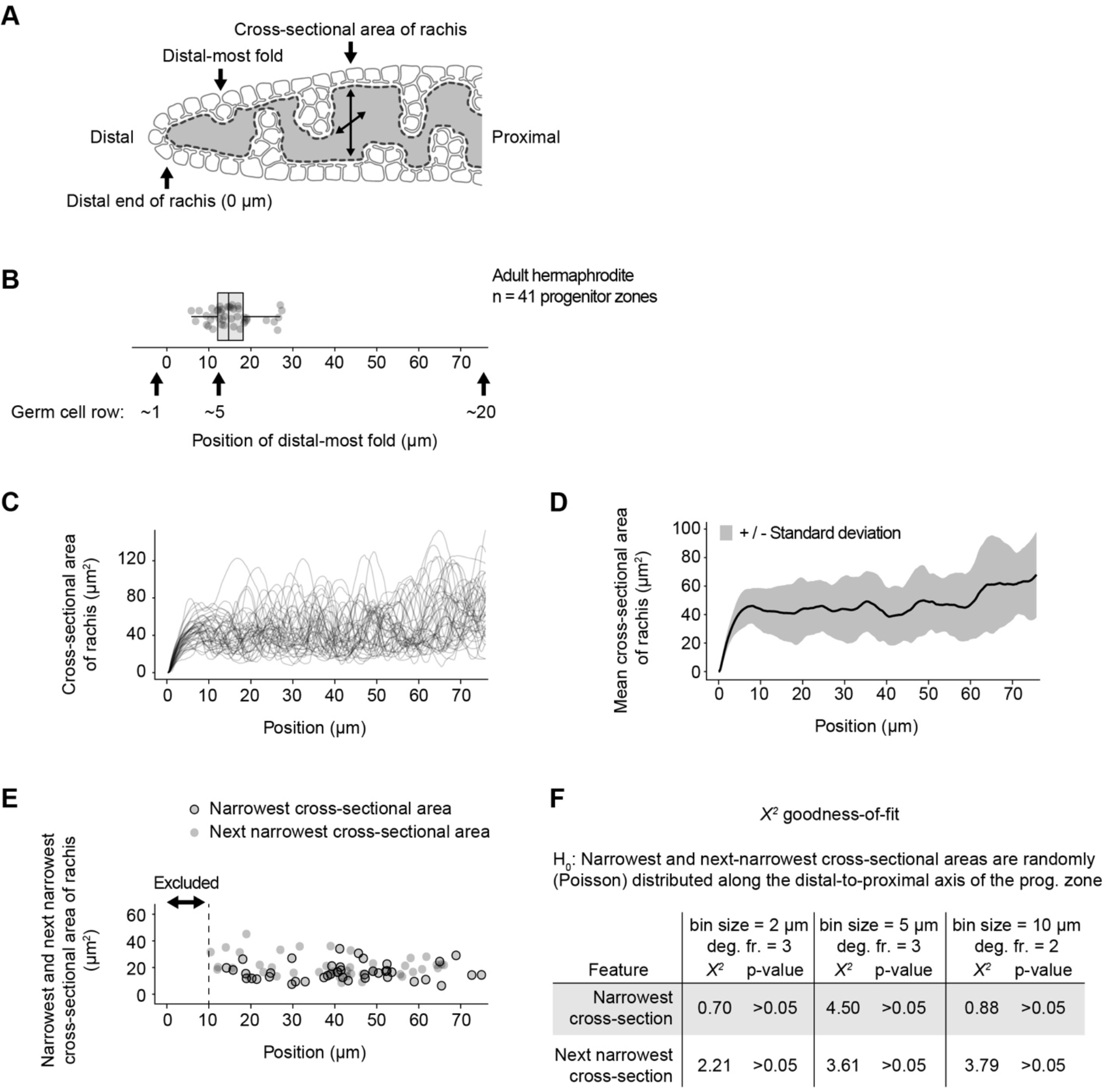
Germline folds are absent from distal-most germ cells but elsewhere are positioned randomly in the progenitor zone. (A-E) Positional analysis of distal-most folds and rachis cross-sectional area in 41 adult hermaphrodite progenitor zones. Cross-sectional area of the rachis provides a read-out of fold position because cross-sectional area is larger where folds are absent or shallow. (A) Schematic of positional analysis. The distal-most tip of the rachis was defined as position 0 μm. (B) Position of distal-most fold. Boxplot: center bar, median; box, interquartile range (IQR); whiskers, most extreme value within 1.5*IQR from box. (C) Cross-sectional area of the rachis. Individual progenitor zones are over-plotted as separate curves. (D) Mean cross-sectional area of the rachis. (E) Narrowest and next-narrowest cross-sectional area of the rachis. This analysis excluded the distal-most 10 μm of the rachis, because the narrowest area necessarily occurs at the distal-most tip. Analysis of next-narrowest area excluded an additional window of 10 μm, centered on the narrowest area, to ensure that the next-narrowest area occurred outside the local minimum of the narrowest area. (F) Results of *X*^*2*^ goodness-of-fit test comparing observed positions of narrowest and next-narrowest areas to positions expected under a random (Poisson) model. Bins of 2 μm, 5 μm, or 10 μm yield Poisson parameter (λ) values of 1.2, 2.9, and 5.6, respectively, therefore spanning the recommended bin sizes for testing against a Poisson model.

**Figure S3.**
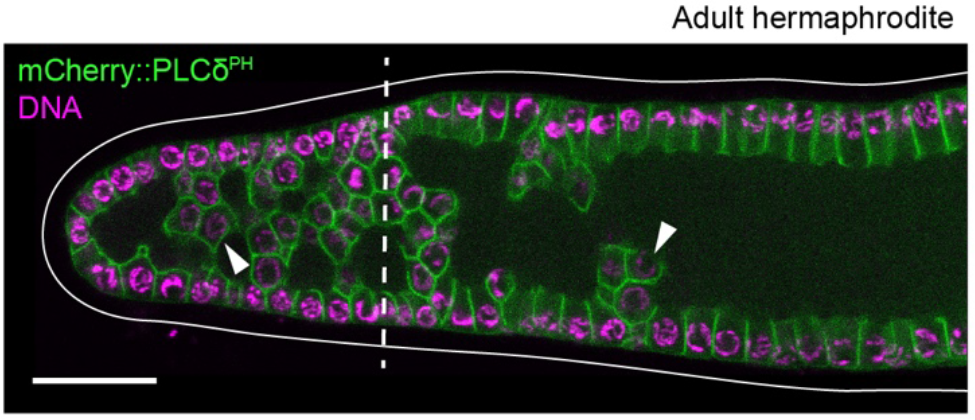
Germline folds visible using the plasma membrane marker mCherry::PLCδ^PH^. Adult hermaphrodite progenitor zone expressing mCherry::PLCδ^PH^ and stained for DNA. Solid line, outline of gonad. Asterisk, distal end of gonad. Dashed line, boundary of progenitor zone. Arrowhead, example of germ cell positioned within a germline fold. Scale bar, 20 μm.

**Figure S4.**
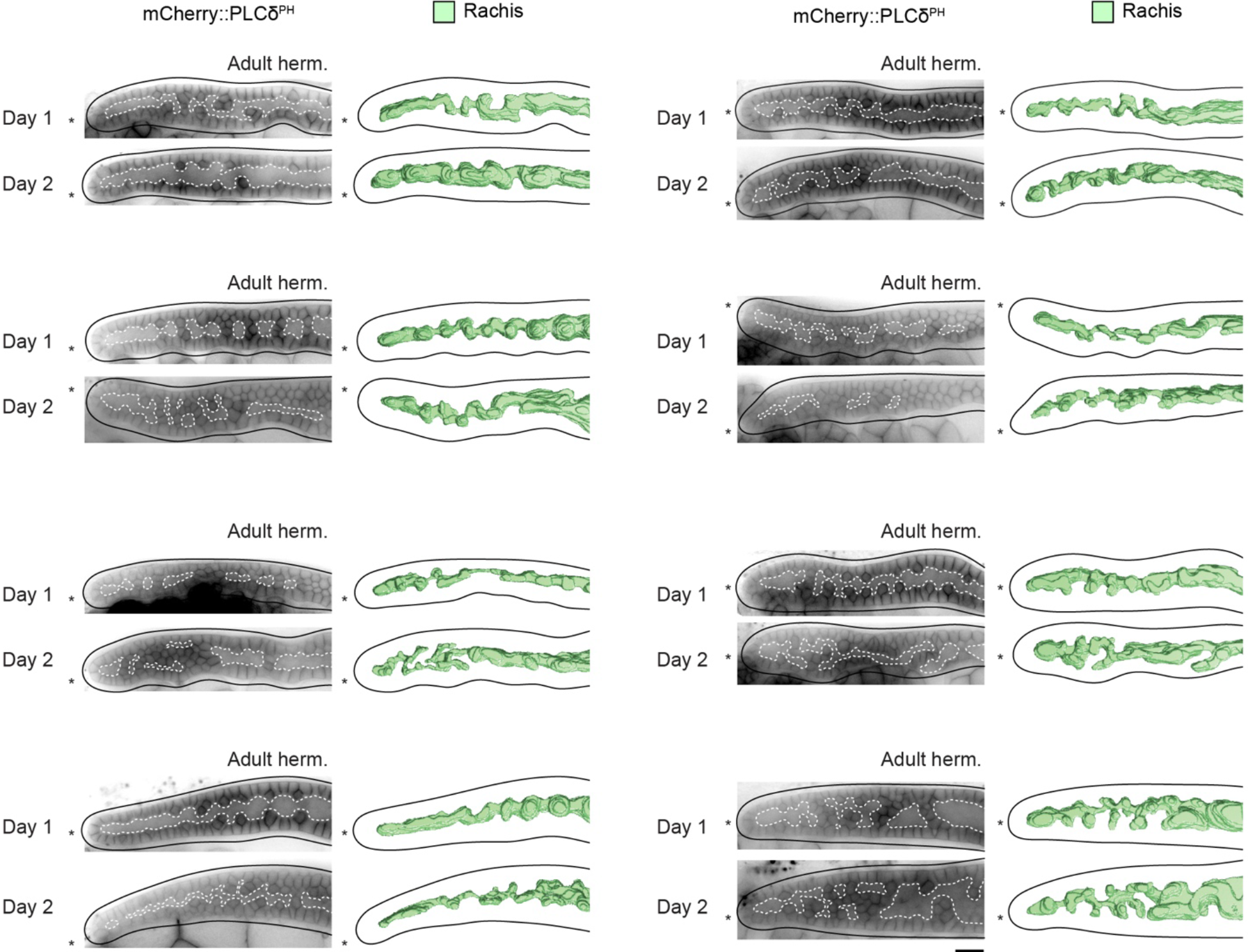
Additional examples of adult hermaphrodite progenitor zones imaged on day 1 and day 2 of adulthood. Left of each pair, adult hermaphrodite progenitor zones expressing mCherry::PLCδ^PH^. Images have been processed with background subtraction. Dashed white line, rachis. Right of each pair, model of rachis. Solid line, outline of gonad. Asterisk, distal end of gonad. Scale bar, 20 μm.

**Figure S5.**
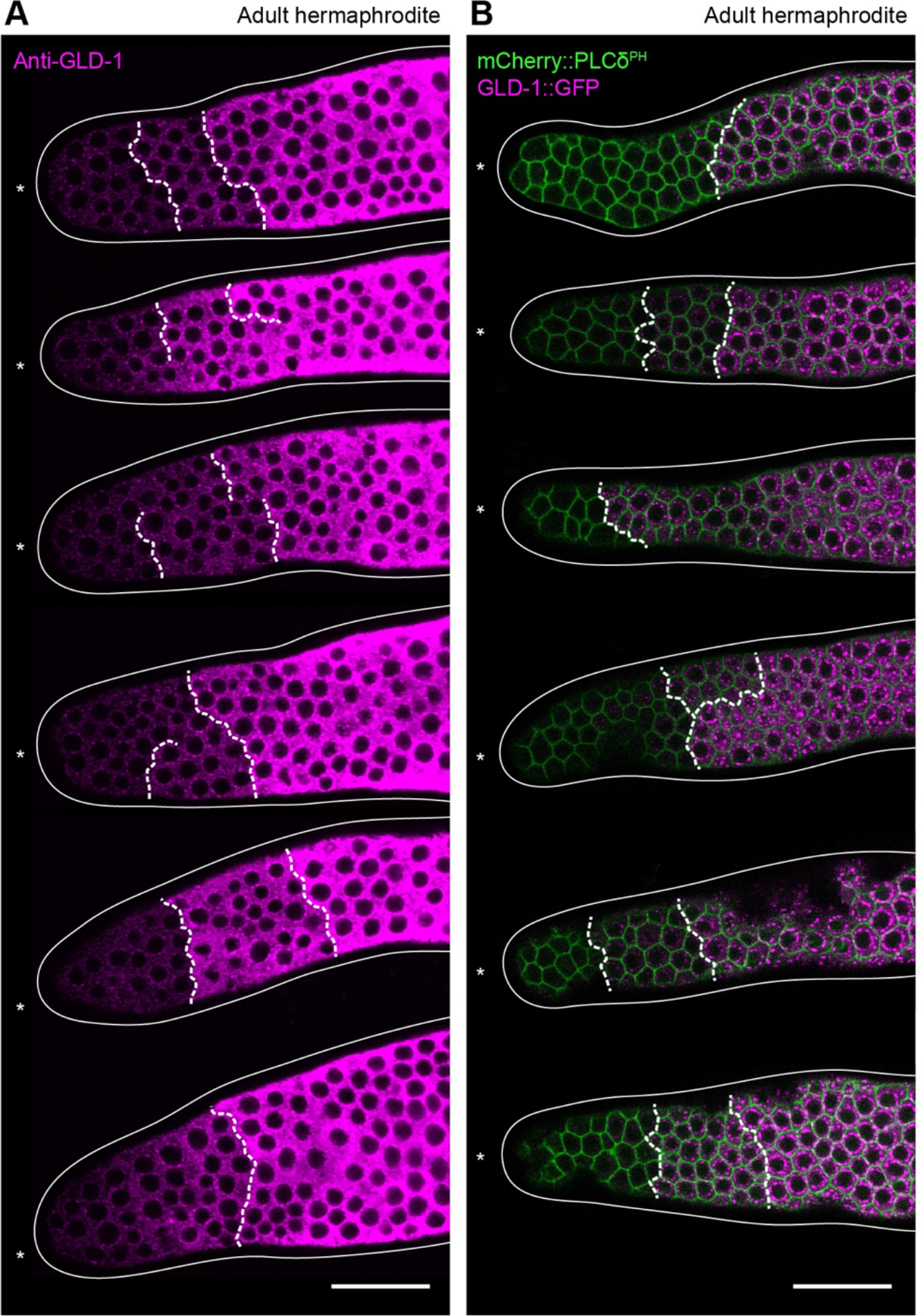
Randomly selected examples of GLD-1 steps. (A) Adult hermaphrodite progenitor zones stained for GLD-1. (B) Adult hermaphrodite progenitor zones expressing GLD-1 ::GFP and mCherry::PLCδ^PH^. (A-B) Solid line, outline of gonad. Asterisk, distal end of gonad. Dashed line, GLD-1 step. Scale bar, 20 μm.

**Figure S6.**
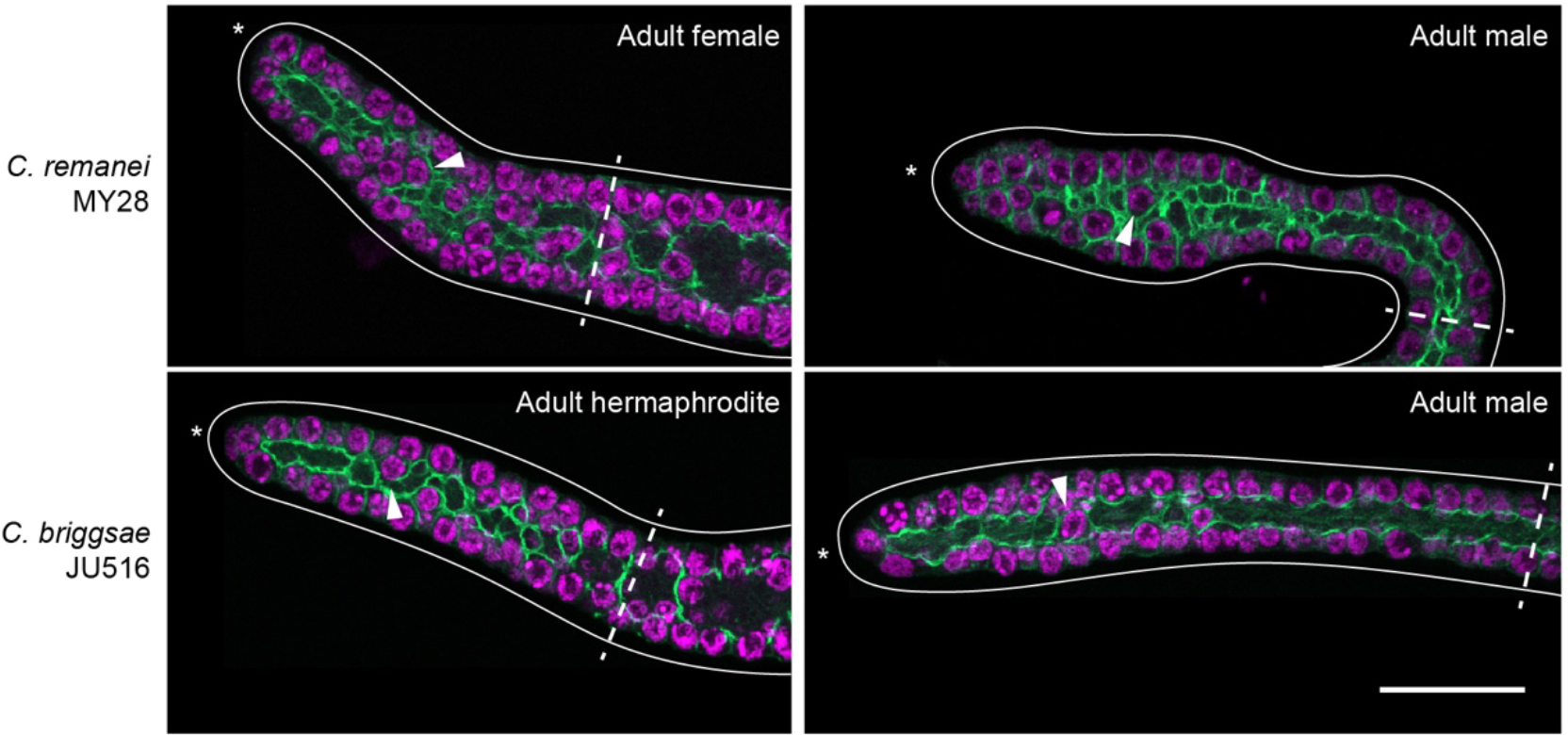
Germline folds in *C. briggsae* and *C. remanei*. Adult *C. briggsae* and *C. remanei* progenitor zones stained for F-actin and DNA. Images are maximum-intensity z-projections through a z-range of 1.5 μm. Arrowhead, example of germ cell positioned within a germline fold. Solid line, outline of gonad. Dashed line, boundary of progenitor zone. Asterisk, distal end of gonad. Scale bar, 20 μm.

